# Actuating tension-loaded DNA clamps drives membrane tubulation

**DOI:** 10.1101/2022.05.02.490361

**Authors:** Longfei Liu, Qiancheng Xiong, Chun Xie, Frederic Pincet, Chenxiang Lin

## Abstract

Membrane dynamics in living organisms can arise from proteins adhering to, assembling on, and exerting force on cell membranes. Programmable synthetic materials, such as self-assembled DNA nanostructures, offer the capability to drive membrane remodeling events in a way that resembles protein-mediated dynamics, but with user-defined outcomes. An example showcasing this capability is the tubular deformation of liposomes by DNA nanostructures with purposely designed shapes, surface modifications, and self-assembling properties. However, stimulus-responsive membrane tubulation mediated by DNA structure reconfiguration remains challenging. Here we present the triggered formation of membrane tubes in response to specific DNA signals that actuate membrane-bound DNA clamps from an open state to various predefined closed states, releasing pre-stored energy to activate membrane deformation. Using giant unilamellar vesicles (GUVs) as a model system, we show that the timing and efficiency of tubulation, as well as the width of membrane tubes, are modulated by the conformational change of DNA clamps, marking a solid step toward spatiotemporal control of membrane dynamics in an artificial system.

## Main

Many cellular processes, such as cell division, vesicular transport and virus infection, involve the tubular deformation of lipid bilayer membranes^1^. Typically, membrane tubulation results from membrane-interacting proteins convening at specific locations in certain orders^2-4^. The well-orchestrated process is coordinated by chemical or mechanical signals responsible for protein recruitment, assembly, disassembly, and conformational change. To better understand the working principle of cells’ arsenal of membrane-deforming machines, a useful practice is to build artificial nanodevices that perform similar tasks on model membranes. Towards this goal, scientists have built a variety of DNA nanostructures that bind lipid bilayers via membrane anchors (e.g., cholesterol and amphipathic peptides)^5-7^. A subset of these nanostructures, designed to mimic BAR, dynamin or the endosomal sorting complex required for transport (ESCRT), can draw membrane tubes from vesicles and supported bilayers^8-10^. The programmable geometry and membrane anchor placement of the DNA nanostructures provide control over parameters that are not readily tunable when working with naturally existing proteins (e.g., membrane affinity, stiffness, self-assembling pattern), thereby shedding light on the determinants of membrane tubulation. However, early examples of membrane-tubulating DNA structures work autonomously, that is, without an ON-switch to the remodeling process after covering membranes with DNA^8,11,12^. Although linker strand and Mg^2+^-mediated DNA-origami polymerization have been shown to induce membrane bulging or tubulation, the membrane remodeling outcomes, particularly the morphology of the deformed membranes, do not necessarily conform to the designed shape of the membrane-coating DNA structures^9,10,13,14^. Moreover, existing membrane-sculpting DNA structures are designed to adopt a single stable conformation, thus lacking the ability to process biochemical signals via conformational changes, a mechanism utilized by proteins like ESCRT-III and dynamin to generate and constrain membrane tubes.

To engineer trigger-responsive DNA devices for better spatiotemporal control of membrane tubulation, we tapped into dynamic DNA nanotechnology, which has developed molecular machines with movable parts and controllable nanoscale motions^15-18^. We are especially interested in a class of mechanical DNA devices, where the bending of a multi-DNA-helix beam can be actuated by the cleavage, folding, or unfolding of single-stranded DNA (ssDNA) domains^19-21^. Based on the hypothesis that bending membrane-anchored DNA nanostructures—specifically their membrane-binding interface—would elicit corresponding curvature changes of lipid bilayers, we built cholesterol-modified DNA clamps containing a prestressed DNA bridge held open by a group of tension-loaded ssDNA strings. Releasing tension via toehold-mediated strand displacement triggered the DNA clamps to close, and in turn led to tubular deformation of liposomes covered by DNA clamps. These DNA clamps thus allowed for on-demand membrane tubulation triggered by specific signals. Interestingly, closing DNA clamps after membrane binding resulted in substantially higher tubulation efficiency than deploying pre-closed DNA clamps to the membrane, possibly because of better membrane-anchor accessibility in the open clamps and simultaneous energy dissipation during the DNA structure actuation. We showed that DNA clamps with different closed states deformed GUVs into tubes with different width, highlighting the programmability of these membrane-sculpting devices.

## Results and discussion

We designed a tension-loaded DNA clamp consisting of two 12-helix-bundle piers (14-nm long each) joined by a 4-helix-bundle bridge (14-nm long) and four ssDNA strings (**Figure 1a**, left). The geometry of an open clamp is codetermined by the gradient of base pair (bp) insertion/deletion (indel) installed in its bridge, which dictates the structure’s curvature at the tension-free state (i.e., closed state)^22^, and the lengths of the tensioned strings, which counteract the effect of the indels and hold the clamp open^23,24^. For example, a bridge with a ±5-bp indel pattern held by four ∼44-nucleotide (nt) strings should theoretically bend slightly (∼18°) in the open clamp, and upon unseating the tensioned (∼10 pN each) strings, close to a higher bending angle of ∼63° (**Figure S1**, also see “Prediction of the bending angle of clamps” in the Supporting Information for details). To facilitate the tension release, we added an 8-nt long overhang to the 5’ end of each ssDNA string, which serves as a toehold to initiate strand displacement when exposed to DNA triggers to detach one end of the strings from the pier (**Figure 1a**, right). The DNA clamp thus stores energy ready to be released by specific DNA triggers in the form of mechanical deformation. Unlike the existing DNA devices that bend in the same direction as the signal-sensing ssDNA domains^20,21^, we purposely placed ssDNA strings on the opposite side of the concave surface so that the DNA triggers have unfettered access to the strings of the membrane-bound clamp.

**Figure 1.**
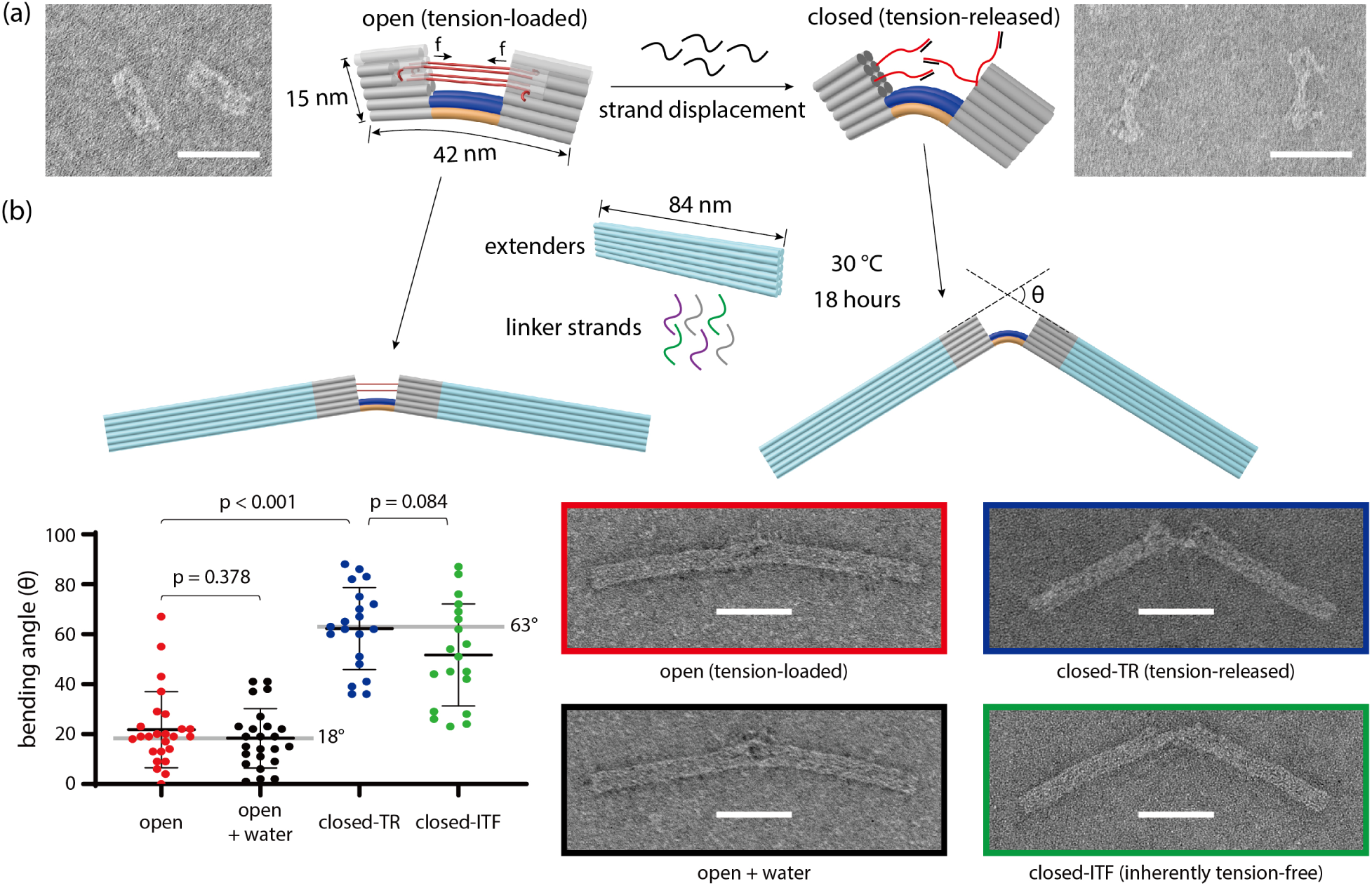
Actuating DNA clamps by triggered strand displacement. (a) Left: An open, tension-loaded clamp consisted of two straight piers (grey cylinders), joint by one bridge (blue/orange cylinders at the bottom) and four ssDNA strings (red lines at the top). The near-flat shape results from the balance between the curved bridge and the tensioned strings. Right: Upon toehold-mediated strand displacement (trigger strands shown in black), one end of the strings is detached from the pier, resulting in a closed, tension-released clamp with increased bending. Negative-stain TEM images are shown next to the cartoon models. (b) Top: Schematics of attaching DNA extenders (cyan) to both ends of the clamp with the help of a set of linker strands. Bottom: bending angle distributions (showing all data points with mean ± SD) of DNA clamps in various states and representative TEM images of extender-attached DNA clamps. Predicted bending angles of the open and closed clamps are noted in the plot for reference. P values are produced by unpaired two-tailed Student’s t-test (two-group comparison). TR: tension-released; ITF: inherently tension-free. Scale bars: 50 nm.

We first prepared the open DNA clamps following a well-established DNA-origami folding and purification pipeline. Briefly, annealing the mixture of a 1512-nt long circular ssDNA (scaffold strand) and a pool of staple strands led to the self-assembly of open clamps, albeit as a minor product (**Figure S2**). To deter the formation of unwanted dimeric structures, we omitted two staple strands in the bridge in the initial annealing and added them back to the folding mixture for a second round of annealing^25^, which helped the correctly folded open clamps become the dominant product (**Figure S3**, see “Design and assembly of DNA origamis” in the Supporting Information for details). Adding DNA triggers to the purified open clamps (trigger:clamp = 40:1, mol/mol) turned them into the closed conformation within 1 hour at room temperature, as shown in the negative-stain transmission electron microscopy (TEM) images (**Figure 1a**).

To allow for better visualization of the DNA clamp structure and reconfiguration, we built two 84-nm-long DNA extenders to attach to both ends of the clamp (**Figure 1b**) via linker strands designed to bridge unpaired scaffold-strand loops at the ends of the DNA-origami structures (**Figure S4 and S5**). As expected, the TEM images of the clamps became much easier to analyze after extender attachment. We measured the bending angles of the open, tension-loaded clamps and closed, tension-released clamps to be 22 ± 15° and 62 ± 16°, respectively, in good agreement with theoretical values. In contrast, adding water to the open clamps did not significantly change their bending angle (18 ± 12°), confirming that the DNA-trigger-mediated strand displacement caused the structural transformation. In addition, we folded clamps without the ssDNA strings (i.e., in an inherently tension-free state, **Figure S1 & S2**), purified them, and measured their bending angle to be 52 ± 20°. We note that the DNA clamps after open-to-close reconfiguration are practically indistinguishable from the inherently tension-free clamps in their bending angle distributions, supporting an efficient structural transformation via the tension-release mechanism.

The stimuli-responsive, flat-to-curved reconfiguration of DNA clamps remotely mimics the conformational change of dynamin, a GTPase responsible for membrane tubule restriction during endocytosis^4^. To enable membrane binding of the DNA clamps, we extended eight ssDNA handles from the concave surface of the clamp for proximal attachment of cholesterol moieties (spaced evenly at ∼5.4 nm) as membrane anchors (depicted as green ellipsoids in **Figure 2a**). Additionally, we labeled the clamp with four copies of Alexa Fluor 647 (depicted as red stars in **Figure 2a**) to facilitate fluorescence microscopy characterization. **Figure 2a** illustrates the experimental procedures for testing the DNA clamp’s membrane binding and deforming activities. Briefly, cholesterol-modified, tension-loaded clamps were mixed with lipid vesicles (99.2 mol % of 1,2-dioleoyl-sn-glycero-3-phosphocholine or DOPC, 0.8 mol % of 1,2-dioleoyl-sn-glycero-3-phosphoethanolamine-N-(lissamine rhodamine B sulfonyl) or Rhod-PE) at a clamp-to-lipid molar ratio of 1:1000, which translates to a theoretical 100% membrane coverage; the mixture was incubated for 1 hour to allow for binding. Subsequently, trigger strands were added; the strand displacement reaction was allowed to run for 1 hour. The entire procedure was carried out at room temperature while keeping osmolarity nearly constant.

**Figure 2.**
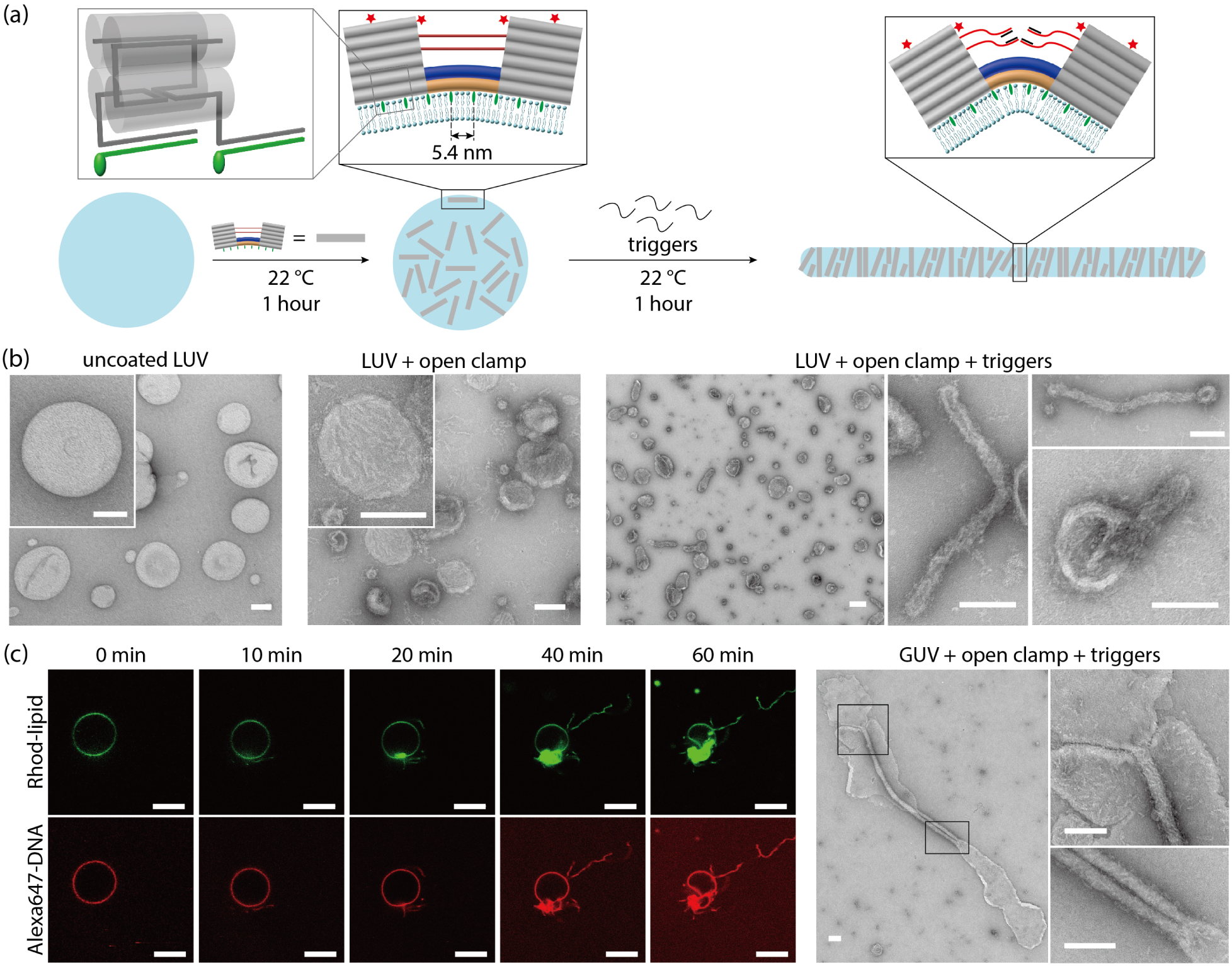
Membrane tubulation driven by DNA clamp actuation. (a) A schematic of vesicle tubulation by actuating membrane-bound DNA clamps. Cholesterol and Alexa Fluor 647 modifications are depicted as green ellipsoids and red stars, respectively. Only staple strands (dark grey lines) of DNA helices (semi-transparent cylinders) are shown in the left cartoon model for clarity. (b) LUV tubulation: TEM images of clamp-free LUVs (left), LUVs coated with open, tension-loaded clamps (middle), and clamp-coated LUVs after triggered conformational change (right). (c) GUV tubulation: a time-course study of tubulation events by confocal fluorescence microscopy (left) and TEM images of tubulated GUVs (right). Scale bars: 100 nm for TEM images and 10 μm for fluorescence microscopy images.

To examine the effect of DNA reconfiguration on membrane morphology, we started with large unilamellar vesicles (LUVs) prepared by lipid-film rehydration and extrusion. After co-incubation with cholesterol-modified DNA clamps in the open conformation, LUVs showed a dense coat of DNA structures under TEM (**Figure 2b**). Most LUVs retained their spherical shape, with only ∼2% appearing to be deformed at this stage. To further evaluate the membrane binding of DNA clamps, the mixture of DNA clamps and LUVs was loaded to the bottom of an iodixanol gradient and spun at 48,000 rpm for 5 hours. Fluorescence scanning and gel electrophoresing the fractions recovered from a post-centrifugation gradient showed that virtually all vesicles co-migrated with DNA to the upper half of the gradient, indicating considerably strong binding between the two, while unbound DNA remained at the bottom (**Figure S6**). Importantly, we observed a surge of tubular structures (diameter: 36.5 ± 9.2 nm, **Table S1**) after treating LUVs covered by open clamps with trigger strands, suggesting membrane tubulation occurred as the result of the conformational change of DNA clamps. Among the deformed vesicles (∼12% of all LUVs), some retained a spherical body with outward protrusions while others turned entirely into a tube. All membrane tubes were covered by DNA structures (**Figure 2b**). To study the influence of the membrane coverage by DNA clamps on vesicle tubulation efficiency, we varied the DNA clamp concentration to achieve theoretical membrane coverage of 0%, 50%, 100%, and 125%. Upon releasing the tension of the DNA clamps, the vesicle tubulation efficiency, defined as the portion of vesicles displaying tubular structures among all vesicles >100 nm in diameter, positively correlated with the initial DNA clamp concentration (**Figure S7**). Tubulation was not detected on vesicles free of DNA clamps. At 50% surface coverage, membrane tubes formed on only 3.0% of LUVs (N = 201). Increasing the coverage to 100% enhanced tubulation efficiency nearly fourfold, reaching 11.9% (N = 67). Further increasing the DNA clamp concentration resulted in a modest increase in DNA tube abundance (efficiency =17.7% at 125% coverage, N = 333). These results are consistent with the concentration-dependent membrane remodeling effects of DNA structures, further supporting the role of the structure-switching DNA clamp in driving vesicle tubulation^8,9^.

To capture the DNA-mediated membrane dynamics in real-time, we next used giant unilamellar vesicles (GUVs, prepared by electroformation) as model membranes. After 1-hour co-incubation of open DNA clamps with pre-adsorbed GUVs on a glass slide, trigger strands were introduced to initiate the strand displacement (defined as time 0). We monitored the membrane tubulation on GUV membranes using confocal fluorescence microscopy for 1 hour (**Figure 2c**, left; **Movie S1**&**S2**). In two time-course studies, outward membrane tubulation became visible at ∼10 min. In the next 10–30 minutes, tubules grew in length and quantity. Continued incubation for up to 1 hour led to further extension of membrane tubules and distortion of the GUV body. Furthermore, the tubular structures showed both rhodamine (from lipid) and Alexa Fluor 647 (from DNA) fluorescence, suggesting the membrane tubes were wrapped by DNA clamps, which was corroborated by TEM imaging of tubulated GUVs (**Figure 2c**, right). These tubes were wider (diameter = 62.1 ± 10.3 nm) than those originated from LUVs, presumably because of a larger lipid reservoir and lower membrane tension of GUV. After 1 hour of strand displacement, ∼61.5% GUVs (N = 39) showed at least one tubular protrusion on the surface (**Figure 3a, 3d**). As expected, we did not detect any tubular deformation on GUVs covered with open clamps, before (N = 38) or after (N = 39) the addition of water in place of the DNA triggers. Surprisingly, when GUVs were incubated with inherently tension-free clamps, membrane tubes appeared on only ∼10.9% of the GUVs (N = 55). In other words, actuating membrane-bound open clamps was about 4× more likely to induce tubulation than directly treating membranes with closed clamps, despite a similar measured surface density of DNA clamps on GUVs (**Figure 3b, 3d**). A similar trend was observed on LUVs (**Figure S26**), although the difference was only ∼2 fold. A possible explanation is that the open conformation of tension-loaded clamps exposed all membrane anchors for near-maximal membrane accessibility, while the closed clamps may obscure cholesterols under the curved bridge, making them less likely to insert into the lipid bilayer. Tubulation thus occurs more readily on vesicles with higher leaflet asymmetry (i.e., more membrane anchors inserted), which promotes spontaneous membrane curvature, and better DNA-membrane contact, which favors curvature coupling between DNA clamp and bilayer. This effect was more prominent on GUVs than LUVs, probably because the near-zero membrane curvature of the former further discriminates against the closed clamps with higher curvature. To test this hypothesis, we built two variants of the inherently closed DNA clamps with only 4 cholesterol moieties per clamp, one with cholesterols attached towards the ends of the structure and the other near the center. Quantifying the Alexa Fluor 647-labeled DNA clamps on the GUV surface showed that the closed clamps labeled with 4 cholesterols near the ends covered GUVs with a surface density comparable to those with 8 cholesterol labels, while the center-labeled variant had significantly lower density on GUVs (**Figure 3b**). Therefore, it is entirely possible for a closed DNA clamp to bind stably with a GUV using only a subset of its membrane anchors, thus unable to utilize the energy generated by membrane insertion of all 8 cholesterols for tubulation. Our data are consistent with the notion that the accessibility of membrane anchors is important for the membrane affinity of DNA nanostructures, and that the energy revenue from membrane-anchor insertion must be sufficient to offset the energy expense of membrane remodeling for membrane-deforming DNA nanostructures^8,26^. However, unlike the static DNA structures, reconfiguring tension-loaded DNA clamps released additional mechanical energy, presumably within a relatively short time window, which contributes towards membrane remodeling.

**Figure 3.**
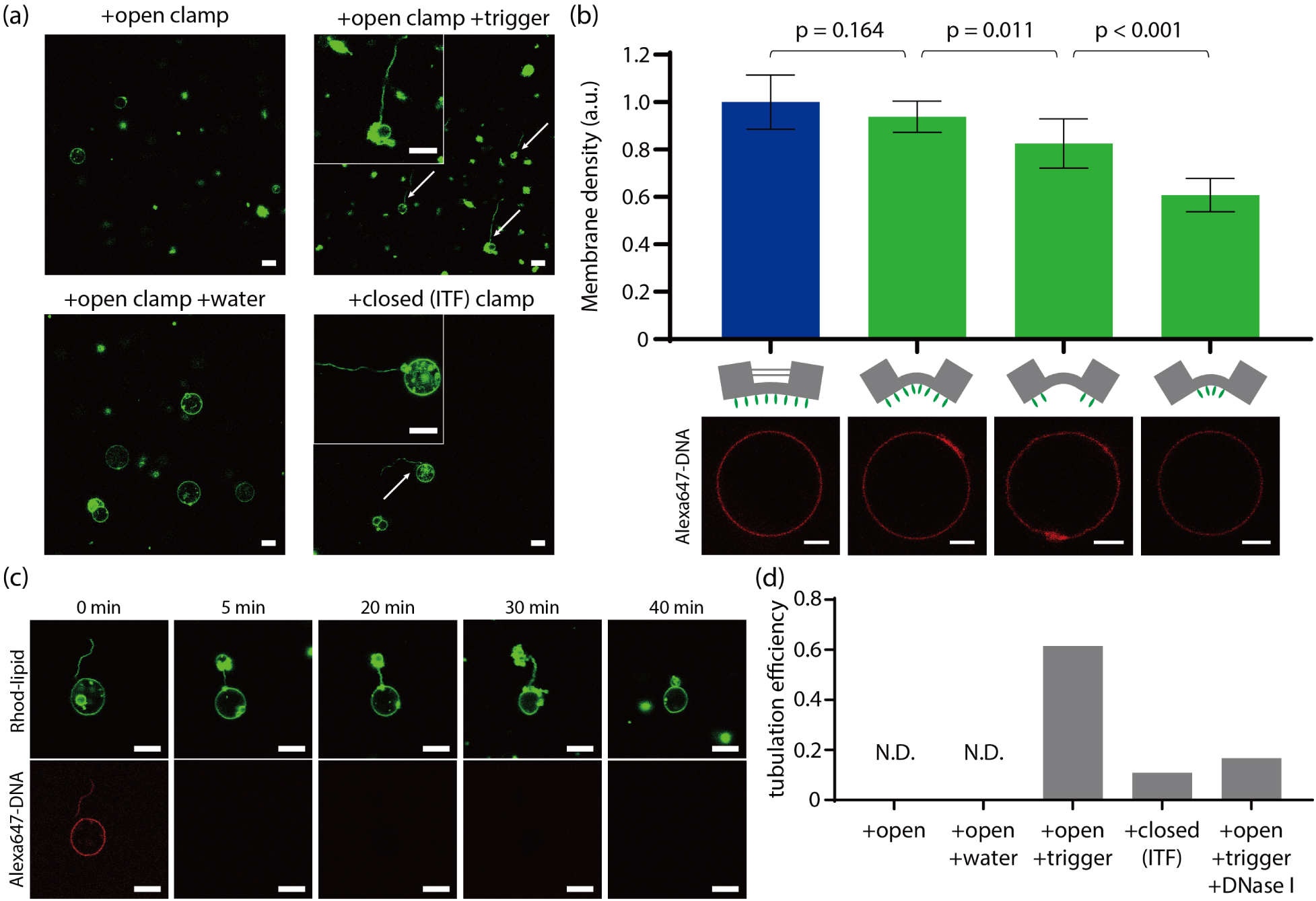
Detailed study of GUV tubulation driven by DNA clamps. (a) GUVs after coincubation with DNA clamps. Reagents added to GUVs are noted on top of the corresponding fluorescence microscopy images (ITF: inherently tension-free). White arrows point to membrane tubes. (b) Quantification of GUV surface coverage by open (blue bar) and closed (green bars) DNA clamps with various cholesterol modifications (schematics shown under the bar graphs). The membrane density of DNA clamps is calculated by dividing the integrated Alexa Fluor-647 (Alexa647) signal (pseudo-colored red in representative microscope images) by the vesicle’s surface area within the confocal volume and normalized to the average density of the open clamp labeled with 8 cholesterols. Bar graphs represent mean ± SD, N=10. P values are produced by unpaired two-tailed Student’s t-test (two-group comparison). (c) Time-course study of a tubulated GUV after DNase I treatment (nuclease added at t=0 min). (d) GUV tubulation efficiency (tubulated GUVs ÷ total GUVs) under various conditions. N.D.: “not detected”. Scale bars: 10 μm.

Membrane tubulating proteins such as dynamin and ESCRT-III are thought to induce membrane scission and vesicle budding by depolymerization and membrane dissociation^27-31^. It is thus interesting to ask whether removing DNA clamps from membranes can mediate the severing of membrane tubes or the formation of budding vesicles. We previously showed that membrane tubes originating from LUVs largely vanished after losing their DNA coat to enzymatic digestion^9^. Here we treated tubulated GUVs with deoxyribonuclease I (or DNase I, an endonuclease that digests both single- and double-stranded DNA) and monitored the membrane dynamics by confocal microscopy (**Figure 3c** and **S8**). Consistent with previous findings, most membrane tubes disappeared 1 hour after the addition of DNase I, suggesting their reliance on DNA coats for stability. A time course study revealed that tubular protrusions from GUVs already started to deform within 5 minutes of nuclease treatment, coincident with the disappearance of Alexa Fluor 647 signals from the GUV surface. At this stage, the membrane tube shortened while remaining connected to its parent GUV, with a second vesicle emerging at the distal end. This asymmetric dumbbell-like structure persisted for as long as 30 minutes, until the tube eventually disappeared, and the distal vesicle departed. In other incidents, the removal of DNA clamps appeared to cause the membrane tubes to break into multiple vesicles (**Figure S8**). It is notable that while most of the fluorescent labels and the cholesterol anchors were cut from the DNA clamps by DNase I within 5 minutes, some partially digested DNA structures existed for up to 40 minutes (**Figure S9**), which may linger and contribute to the tubular membranes that survived longer. Therefore, our data suggested a possible mechanism to artificially induce vesiculation by stripping narrow (tens of nanometers in width) membrane tubes of their stabilizing DNA coats.

Our working model is that the DNA clamps induce membrane tubulation by releasing the energy stored in the prestressed DNA bridge and imposing curvature of the clamps on the membrane. Therefore, a reasonable hypothesis is that the eventual abundance and width of membrane tubes are tied to the amount of energy initially stored in the tension-loaded clamps. To systematically test this, we built a set of reconfigurable DNA clamps (named I, II, III, IV and V) with nearly flat open conformations but with increasing curvatures in the closed conformations (**Figure S10**). All 5 clamps were designed using a common principle (minor changes are noted in the Supporting Information under “Design considerations for highly curved DNA clamps”). Therefore, those storing more energy in the tension-loaded (i.e., open) state should exhibit higher curvatures once the tension is released by triggered strand displacement. Assembling and actuating the DNA clamps in solution confirmed that all 5 versions of DNA clamps folded with a decent yield and underwent structural transformation in response to the trigger strands (**Figure S11**). In general, most clamps adopted curvatures in good agreement with the design before and after reconfiguration, as measured from negative-stain TEM images (**Figure 4a–4b** and **S12–S17**). Notably, clamps IV and V, the two versions storing the most energy, folded with more defects (**Figure S18**) and greater deviation from expected curvatures (**Figure 4b** and **S17**) than the rest of the set. This is not surprising as DNA structures containing severely bent helices or highly tensioned ssDNA segments (f >20 pN) are known to be prone to misfolding^22,32^. Nevertheless, the clamps provided a toolset to investigate the correlation between the mechanical properties of DNA devices and their membrane tubulating functions.

**Figure 4.**
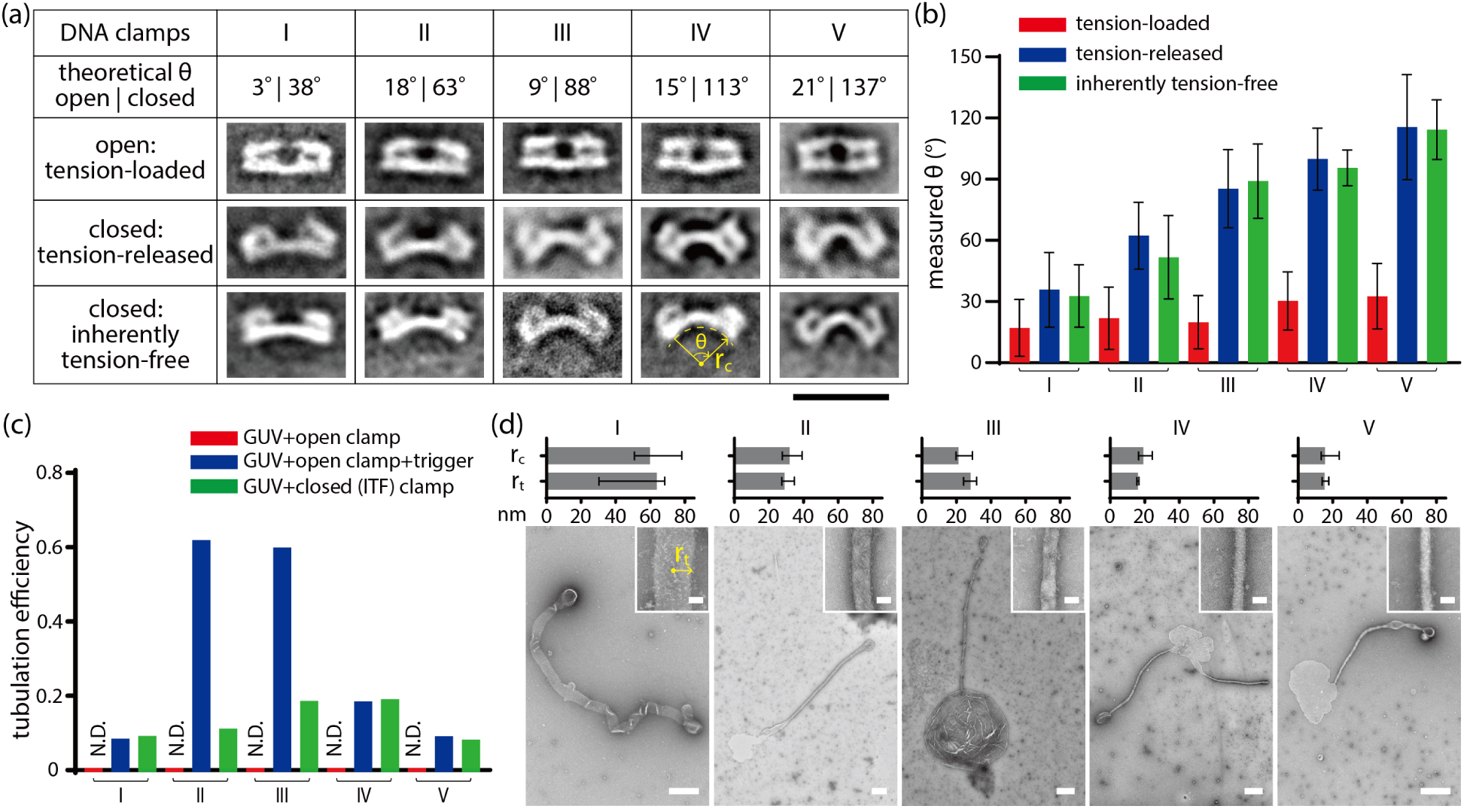
GUV tubulation by DNA clamps with different curvatures. (a) Class-average TEM images of five different DNA clamps (named I–V with increasing curvatures) in open and closed conformations. Scale bar: 50 nm. (b) Bending angles (θ) measured from extender-attached DNA clamps. Bar graphs represent mean ± SD, N=11–28. (c) Efficiency of GUV tubulation induced by DNA clamps. N.D.: not detected; ITF: inherently tension-free. (d) Morphology of membrane tubules covered by DNA clamps with varying curvatures. Bar graphs (top) show the medians of r_c_ (radii of curvature of closed DNA clamps) and r_t_ (radii of membrane tubes) with 95% confidence intervals. TEM images (bottom) show representative tubulated GUVs (scale bars: 400 nm). Insets show magnified regions of membrane tubes (scale bars: 50 nm).

When deployed to GUVs, all 5 DNA clamps generated membrane tubes in a trigger-dependent manner (**Figure 4c** and **S19**). Actuating tension-loaded DNA clamps on membrane generally resulted in similar or higher tubulation efficiency compared with covering GUVs with inherently tension-free clamps. These behaviors are well represented by clamp II that we initially tested. Unsurprisingly, clamps II and III drew much more (∼6×) tubes from GUVs than clamp I, which probably does not release enough energy to nucleate the formation of a membrane tube. However, the tubulation efficiency did not increase further with the higher energy-storing capabilities of clamps IV and V, which showed considerably diminished membrane remodeling activities. We attribute this to the relatively high occurrence of structural defects in these highly tensioned clamps (**Figure S18**). Such defective clamps can still bind to GUVs but may be unable to close and disperse energy properly, thus impeding the membrane tube elongation (see “Proposed mechanism of membrane tubulation” in the Supporting Information for details). Interestingly, clamps IV and V tubulated LUVs as efficiently as clamps II and III (**Figure S26**), suggesting that some misfolded clamps may still participate in membrane remodeling, but their involvement makes it difficult to form membrane tubes of sufficient length to be detected by fluorescence microscopy.

We next examined how the curvature of closed DNA clamps might influence that of the membrane tubes. For this, we calculated the radii of curvature (r_c_) of the tension-released DNA clamps from their measured bending angles (**Figures S17** and **S20**). The r_c_ of each DNA clamp was then compared with the radius of membrane tubes (r_t_) generated by actuating the clamp on GUVs (**Figure 4d**). Strikingly, we found nearly identical r_c_ and r_t_ values for the entire set of clamps, strongly suggesting that the tension-release mechanism bent DNA clamps on GUVs as much as in solution and that the closed clamps wrapped membranes tightly following the circumference of the tubes (**Figure S20**). However, this trend does not hold for LUVs (**Figure S21-25, Table S1**). Despite their very different shapes in the closed state (r_c_ averaging 15–60 nm), all five DNA clamps gave rise to membrane tubes of similar widths (mean r_t_ = 15–20 nm). Thus, there seemed to be other determinants in addition to the geometry of DNA clamps that defined the width of membrane tubes. We suspect the small size of extruded liposomes limited the lipid supply and led to a steep rise in membrane tension during membrane tubulation, making it difficult to form wide tubes of appreciable length (see “Proposed mechanism of membrane tubulation” in the Supporting Information for details).

## Conclusions

DNA nanostructures with curved membrane-binding interfaces have shown their promise as programmable membrane-remodeling tools^8-10^. However, to fully recapitulate the well-regulated subcellular membrane dynamics in an artificial system, there is a pressing need for signal-responsive nanodevices that manipulate membranes with predictable outcomes. The tension-loaded DNA clamps presented here stably bind to membrane in an inactive form and transform into the remodeling-competent form only when DNA triggers release their internal tension, thereby providing a means to activate membrane-remodeling nanodevices with specific biochemical signals. Consistent with our design, we show that the DNA clamps tubulate GUVs most efficiently when the DNA structural transformation on the membrane releases energy sufficient for tube nucleation and elongation. Moreover, the width of GUV-originated membrane tubes is dictated by the curvature of the DNA clamp’s cholesterol-labeled surface, offering the opportunity to control membrane topography with rationally designed DNA nanostructures. We envision future development in the following areas. First, although our proof-of-concept study has shown the design principle can be generalized to build dynamic DNA structures with various geometrical and mechanical properties, DNA clamps with high internal stress fold with suboptimal quality which negatively impacts their membrane tubulation efficiency. Alternative design and assembly methods that improve the integrity of the prestressed DNA nanostructures are thus desirable. Second, the DNA clamps are strong enough to deform synthetic vesicles ranging from several hundred nanometers to tens of micrometers in diameter. It would be interesting to see how such DNA devices perform on the plasma membrane of cells with complex chemical composition and underlying cytoskeleton. Third, with a rich library of nucleic acid chemistry and well-developed DNA-based logic gates^33-40^, it should be possible to build membrane-deforming devices with sophisticated control mechanisms and diverse molecular triggers. Finally, the DNA clamp’s ability to recognize and process DNA signals opens opportunities to recruit and coordinate nanodevices by messenger molecules, such that devices with different functions can work in concert to accomplish complicated tasks, such as sorting membrane-associated cargos, packaging them into vesicles, and delivering them to designated locations.

## Acknowledgment

We thank the Yale West Campus Imaging Core for the assistance with fluorescence microscopy. This work is supported by National Institutes of Health grants GM132114 and GM141669 and a Yale University faculty startup fund to C.L., an Agency for Science, Technology and Research Graduate Scholarship (Singapore) to Q.X., a China Scholarship Council fellowship to C.X.

## Author contributions

L.L. initiated the project, designed and performed most of the experiments, analyzed the data, and prepared the manuscript. Q.X. assisted with vesicle preparation. C.X. prepared p1512 scaffold strand for folding DNA clamps. F.P. interpreted the data, proposed the mechanism of tubulation, and prepared the manuscript. C.L. initiated the project, designed and supervised the study, interpreted the data, and prepared the manuscript. All authors reviewed and approved the manuscript.

## Supporting Information Available

Materials and methods, notes, as well as supplemental figures, tables, and videos, are available in the supporting information.

## Data availability

All raw data associated with this study are available from the corresponding authors upon reasonable request.

