## Supplementary Information for "Actuating tension-loaded DNA clamps drives membrane tubulation"

### TABLE OF CONTENTS

MATERIALS AND METHODS ... 3
Design and assembly of DNA-origami structures ... 3
Preparation of large unilamellar vesicles (LUVs) ... 4
Preparation of giant unilamellar vesicles (GUVs) ... 4
Transmission electron microscopy (TEM) ... 4
Confocal fluorescence microscopy ... 4
DNA-membrane affinity examined by density gradient centrifugation ... 5
Agarose gel electrophoresis ... 5
NOTES ... 5
Design considerations for highly curved DNA clamps ... 5
Prediction of the bending angle of clamps ... 6
Calculations for closed clamps ... 6
Calculations for open, tension-loaded clamps ... 7
Proposed mechanism of membrane tubulation ... 12
Energetics of tube formation ... 13
Nucleation of a tube ... 14
Elongation of a tube ... 14
Tension-driven tube diameter ... 15
SUPPLEMENTAL FIGURES ... 17
Figure S1. ... 17
Figure S2. ... 18
Figure S3. ... 19
Figure S4. ... 20
Figure S5. ... 21
Figure S6. ... 22
Figure S7. ... 23
Figure S8. ... 24
Figure S9. ... 25
Figure S10. ... 26
Figure S11. ... 27
Figure S12. ... 28
Figure S13. ... 29
Figure S14. ... 30

Figure S15. 31
Figure S16. 32
Figure S17. 33
Figure S18. 34
Figure S19. 35
Figure S20. 36
Figure S21. 37
Figure S22. 38
Figure S23. 39
Figure S24. 40
Figure S25. 41
Figure S26. 42
SUPPLEMENTAL TABLES 43
Table S1. 43
Table S2. 44
REFERENCES 47

### MATERIALS AND METHODS

#### Design and assembly of DNA-origami structures

The DNA clamps and extenders were designed in caDNAno (**Figure S1, S4, S10, and S11**)[1]. All DNA oligonucleotides were purchased from Integrated DNA Technologies. Unmodified staple strands (excluding the ssDNA strings) were purchased in 96-well plate format with concentrations normalized to 100 µM. Oligonucleotides longer than 80 nt (ssDNA strings) were purchased in tube format and purified in house by denaturing PAGE. Oligonucleotides with fluorescent or cholesterol modifications (anti-handles) were HPLC-purified by the vender. Scaffold strands p1512 and p3024 were prepared using a previously reported method[2-4].

The open, tension-loaded clamps (**Figure 1a**, left **and S1**) were assembled in two steps. Step 1: Mix p1512 scaffold strand (50 nM) and a pool of staple strands (300 nM each), excluding two staple strands in the bridge (inside the black box in **Figure S1**), in a TE buffer (5 mM Tris-HCl, 1 mM EDTA, pH 8.0) containing 12.5 mM $MgCl_2$, using a 36-hour thermal annealing protocol (80−65 °C, -1 °C/5 min; 64−24 °C, -1 °C/50 min; 12 °C hold). Step 2: Add the two staples back into the solution above and anneal the batch using an 18-hour protocol (40−20 °C, -0.1 °C/5 min; 12 °C hold). The two-step folding strategy is deemed to promote the folding of properly folded clamps, likely because incorporating ssDNA strings into the piers before forming the bridge reduces the clamp oligomerization mediated by ssDNA strings (**Figure S2**–**S3**).

The closed, inherently tension-free clamps (**Figure S1**) were assembled from p1512 scaffold strand (50 nM) and a pool of staple strands (300 nM each), excluding the four ssDNA strings, in a TE buffer containing 12.5 mM $MgCl_2$, using the 36-hour protocol.

The two extenders (**Figure 1b and S4**) were assembled from p3024 scaffold strand (50 nM) and a pool of staple strands (300 nM) in a TE buffer containing 10 mM $MgCl_2$, using the 36-hour protocol.

All folded DNA origami structures, except the open, tension-loaded clamp V, were purified by PEG (polyethylene glycol) fractionation[5,6]. Briefly, the assembled structures were supplemented with 8% w/v PEG-8000 and 0.5 M NaCl and held at room temperature for 10 min before centrifugation at 15000 g, 4 °C for 15 min. The post-centrifugation supernatant was carefully removed by pipetting. The pellet was re-suspended in a TE buffer containing 12.5 mM $MgCl_2$, followed by another round of PEG fractionation. The pellet was finally dissolved in a TE buffer containing 12.5 mM $MgCl_2$ and stored at 4°C before usage.

The open, tension-loaded clamp V was purified by rate-zonal centrifugation[7]. Typically, 0.5 mL of assembled product was concentrated to 0.2 mL using PEG fractionation, loaded on top of a 15−45% (v/v) quasi-linear glycerol gradient in a polycarbonate centrifuge tube (13×51 mm, Beckman Coulter Inc.), and spun at 50,000 rpm in a Beckman swing bucket rotor (SW55-Ti rotor) for 3 hours. The contents of the tube were fractionated from top to bottom (200 µL per fraction). Eight microliter of each fraction was then loaded onto a 1.5% agarose gel containing 0.5 µg/mL ethidium bromide (EtBr) and run in 0.5× TBE (45 mM Tris-Base, 45 mM boric acid, and 1 mM EDTA), 10 mM $MgCl_2$ for 2 hours at 5 V/cm. After image analysis on a Typhoon FLA 9500 imager (GE Healthcare), the fractions containing well-formed monomeric DNA clamp were combined and concentrated using PEG fractionation. The pellet was then dissolved in a TE buffer containing 12.5 mM $MgCl_2$ and stored at 4°C before usage.

The concentrations of DNA origami structures were determined using a NanoDrop 2000 (Thermo Fisher).

To attach DNA extenders to the clamp, a mixture of DNA clamp (5 nM), extenders (6 nM each), and linker strands (100 nM each) in a TE buffer containing 12.5 mM $MgCl_2$ were incubated at 30 °C for 18 hours.

#### Preparation of large unilamellar vesicles (LUVs)

LUVs composed of 99.2% DOPC and 0.8% Rhod-PE (lipids purchased from Avanti Polar Lipids) were produced by lipid-film rehydration and extrusion. Briefly, appropriate amounts of lipids in chloroform were mixed in glass tubes and dried in nitrogen gas for 30 min and under vacuum overnight. The lipid film was then suspended in 300 µL of hydration buffer (25 mM HEPES-KOH, 100 mM KCl, 10 mM $MgCl_2$, pH 7.4) by agitation to achieve a final lipid concentration of 1 mM. The suspended lipids were frozen-thawed in plastic centrifuge tubes for seven cycles, each consisting of 15-s flash-freezing in liquid nitrogen followed by 2-min water-bathing at 37 °C. Final homogenization was achieved through 40 forward-and-back extrusion pumps using an Avanti mini-extruder with a 200-nm filter. The final LUVs were stored at 4 °C for no more than 2 weeks before usage.

#### Preparation of giant unilamellar vesicles (GUVs)

GUVs composed of 99.2% DOPC and 0.8% Rhod-PE were produced by electroformation[8]. Briefly, to form thin lipid films on ITO glass slides, 125 µL of 1 mg/mL lipid solution in chloroform was spotted on the conducting face of two glass slides within a marked 3 cm × 3 cm area; the chloroform was then evaporated on a 50 °C heating plate. One piece of copper tape was placed on the conducting face of each slide, extending over the edge. Two of such ITO slides with lipid-film-covered areas were aligned over each other, separated by a silicon slab containing a 3 cm × 3 cm × 3 mm (L×W×H) hole inside, to create a chamber accessible only via a syringe entry channel with copper tape extensions on opposite sides of the chamber. The sandwich was held together with binder clips, wrapped in foil, and placed in a vacuum chamber. After 1 hour, the chamber was filled with about 3 mL of sucrose solution containing 0.03% w/v sodium azide at an osmolarity of 207 mOsm (measured via a Thomas Scientific Micro-Osmette osmometer), which is 16 mOsm lower than the hydration buffer (223 mOsm) to prevent vesicle bursting. The filled chamber was sheltered from light with aluminum foil. The copper strips were connected to a waveform generator initially set at a frequency of 10 Hz, 0 phase, and 100 mVpp amplitude. The amplitude was gradually increased every 6 minutes as follows: 200 mV, 300 mV, 500 mV, 700 mV, 900 mV (each at 10 Hz), and then left at 1.2 V at 10 Hz for 1 hour, finishing with 1.4 V at 4 Hz for 30 min. The resulting GUVs were then carefully extracted using a 1.1 mm needle on a 2 mL syringe and stored in LoBind Eppendorf tubes at 4 °C.

#### Transmission electron microscopy (TEM)

For DNA-only samples, a drop of the sample (5 µL) was deposited on a glow discharged formvar/carbon-coated copper grid (Electron Microscopy Sciences), incubated for 1 min, and blotted away. The grid was first rinsed twice with 5 µL of TE buffer containing 12.5 mM $MgCl_2$, washed briefly with 5 µL of 2% (w/v) uranyl formate, and stained for 1 min with 5 µL of 2% uranyl formate. For samples containing liposomes, a drop of the sample (5 µL) was deposited on a glow discharged formvar/carbon-coated copper grid, incubated for 4 min, and blotted away. The grid was then rinsed with 2% uranyl formate for 10 s and stained with 2% uranyl formate for 1 min. TEM images were acquired on a JEOL JEM-1400Plus microscope (acceleration voltage: 80 kV) with a bottom-mount 4k×3k CCD camera (Advanced Microscopy Technologies). Negative stain 2D class averages were computed using EMAN2[9].

#### Confocal fluorescence microscopy

A 10 µL drop of 2 mg/mL bovine serum albumin (BSA) was placed on an uncoated MatTek glass-bottom dish, let sit for 20 min, and washed with hydration buffer. After that, a 10 µL drop of diluted GUV stock (~1.6× volume of hydration buffer added to 1 volume of stock) was dispensed on the BSA-coated region of the glass-bottom dish and let sit for 20 min to allow for GUV sedimentation on the glass surface. Subsequently, the unbound GUVs were removed by careful pipetting, immediately followed by the addition of 10 µL of hydration buffer (for imaging uncoated GUVs) or Alexa Fluor 647 and cholesterol-labeled DNA clamps (20.4 nM) in hydration buffer (for imaging DNA-coated GUVs). The mixtures were incubated for 1 hour. To actuate DNA clamps for membrane tubulation, 0.33 µL of trigger strands (25 µM each, 40-

fold excess) was added to clamp-coated GUVs and incubated for 1 hour. Alternatively, 0.33 µL of water was added as a negative control. To remove DNA coat from GUVs, 0.5 µL of DNase I stock solution (1 U/µL, Thermo Scientific) was added to tubulated GUVs and incubated for 1 hour. The GUVs were imaged under a Leica TCS SP8 confocal microscope using an HC PL APO CS2 63× OIL objective lens. All experiments were conducted at room temperature. One milliliter of hydration buffer was spotted along the dish walls to mitigate evaporation. For transferring GUVs, 20 µL micropipette tips were cut transversely to increase tip entry diameter and reduce shear stress on GUVs. The clamp density ($c$) on the GUV surface was determined by the comparison between the Alexa Fluor 647 fluorescence density from the GUV surface and that in the bulk solution.

##### DNA-membrane affinity examined by density gradient centrifugation

Alexa Fluor 647 and cholesterol-modified DNA clamp (20.4 nM) were mixed with Rhod-labeled LUVs (20 µM lipid) in 60 µL hydration buffer. This solution was incubated for 1 hour at room temperature for binding and then mixed with 120 µL of 45% iodixanol (STEMCELL Technologies) in hydration buffer. The 180 µL solution (30% iodixanol) was added to the bottom of a 0.8-ml ultracentrifuge tube. Six additional 80-µL layers of iodixanol solution in hydration buffer (26%, 22%, 18%, 14%, 10%, and 6% iodixanol per layer) were carefully stacked on top to form a final 6−30% iodixanol gradient. Gradients were spun at 48,000 rpm at 4°C in a SW55-Ti rotor for 5 hours. Fractions (42 µL each) were then collected from top to bottom of the gradient into a 96-well plate and imaged on a Typhoon FLA 9500 scanner for Alexa Fluor 647 and Rhod fluorescence, either directly or after agarose gel electrophoresis (see below).

##### Agarose gel electrophoresis

Samples and 1 kb DNA ladder (New England Biolabs) were loaded into separate wells of a 1.5% or 2% agarose gel casted in running buffer (0.5× TBE, 10 mM $MgCl_2$) with 0.5 µg/mL EtBr and run for 2 hours at 5 V/cm in running buffer. For lipid-containing samples, a final 0.05% sodium dodecyl sulfate (SDS) was added to running buffer. The gel was scanned on a Typhoon FLA 9500 scanner. To recover purified DNA structures, bands of interest were excised on a UV transillauminator (VWR International) using a razor blade and spun in Freeze ’N Squeeze DNA gel extraction spin columns (Bio-Rad Laboratories).

#### NOTES

##### Design considerations for highly curved DNA clamps

The curvature of closed DNA clamps is solely determined by the gradient of bp insertion/deletion (indel) installed in their bridges. In the open, tension-loaded DNA clamps, ssDNA strings pull against the bent bridge, resulting in a near-flat conformation (see **Prediction of the bending angle of clamps** for more details). Although clamps I, II and III folded with good yield, clamps IV and V, the two versions storing the most energy, folded with undesired multimers (**Figure S11**). One possible explanation is that staple strands hybridizing to the scaffold strand with only ~8-bp uninterrupted segments (boxed in **Figure S11**, left) are unable to sustain such high strains and partially detaches from the clamp to form aggregates with other defective clamps. To mitigate this problem, we designed a new set of staple strands for the highly stressed bridges of clamps IV and V, named “step 2 rev”, by removing a crossover (boxed in **Figure S11**, right) to allow ~16-bp continuous staple-scaffold hybridization. Indeed, the revised design enhanced the yield of correctly folded clamp IV and V; byproducts (dimers and aggregates) were greatly reduced from 74% to a maximum of 34% (clamp V, determined by densitometry of agarose gel images). We therefore assembled clamps IV and V using the “step 2 rev” staple strands for subsequent experiments.

**Prediction of the bending angle of clamps**

The energy stored in the bridge at different bending angles[10] and that in the tensioned ssDNA strings[11] were calculated. The theoretical equilibrium bending angle for the open, tension-loaded clamps was then determined by minimizing the total energy.

Calculations for closed clamps

The energy stored in the bridge was calculated according to the toy model[10]. Briefly, it was calculated by a sum of the bending and stretch/compression energy stored in each helix of the bridge, using the following functions:

$$E_{bridge} = \sum_{i=0}^{3}\left[\frac{1}{2}\cdot S\cdot n_i\cdot\frac{\left(d_i-d_{eq}\right)^2}{d_{eq}}+\frac{1}{2}\cdot B\cdot n_i\cdot\frac{d_{eq}}{(r_0+\delta_i)^2}+\frac{1}{2}\cdot D\cdot\Phi_i\cdot\left(\frac{d_i}{d_{eq}}-1\right)\right]$$

$$d_i = \frac{n_0}{n_i}\cdot d_0\cdot\left(\frac{\delta_i}{r_0}+1\right)$$

$$\Phi_i = 2\cdot\pi\cdot\frac{N-n_i}{10.5}$$

where *S*, *B*, and *D* denote the stretch, bending, and twist-stretch-coupling moduli of DNA, with a set of values (*S* = 660 pN, *B* = 230 pN·nm$^2$, *S* = 0 pN·nm) calculated *via* Young's modulus.

As an example, the following Python (v 3.7.3) script calculates the energy stored in a 4-helix-bundle bridge with indel of ±3 bp at integer bending angles between 1° and 180° and outputs the predicted bending angle with the minimal total energy.

```
# Calculations for inherently tension-free clamps

from numpy import pi, sqrt

# All constants are listed below

D = 2.25E-9 # Diameter of dsDNA (m)
Lds = 0.335E-9 # length of dsDNA (m/base)
kb = 1.38E-23 #Boltzmann constant (J/K)
T = 298 # Temperature(K)
S = 660E-12 # Stretch moduli of DNA helix (N)
B = 230E-30 # Bending moduli of DNA helix (N*m2)


# All adjustable parameters are listed below

Nds = 4 # number of helices in the bridge
Nbp = 42 # number of bp in the bridge without insertion or deletion
L_conn = Nbp*Lds # length of the bridge
indelbp = 3 # bp(n) is the number of bp inserted or deleted from each outer or inner layer of
helices (Note: n can be 3, 5, 7, 9, or 11)
delta_coefficient = 1.25E-9 # each DNA helix is modeled as a 2.5 nm wide rod

n = [0 for i in range(Nds)] # n is the number of bp installed in helices
n[0] = Nbp + indelbp
n[1] = Nbp + indelbp
n[2] = Nbp - indelbp
n[3] = Nbp - indelbp

delta = [0 for i in range(Nds)] # distance of a helix from the bending axis (nm)
delta[0]  = 1.0
```

```
delta[1]  = 1.0
delta[2]  = -1.0
delta[3]  = -1.0

for i in range(Nds):
         delta[i] *= delta_coefficient


# Calculate the energy stored in the bridge to find the bending angle with the energy minimum

helix_total_energy = []
for angle_in_degrees in range(1,180):
        angle_in_radians = angle_in_degrees/180.0*pi
        rref = L_conn /angle_in_radians
        helix_stretch_energy = 0
        helix_bending_energy = 0
        for i in range(Nds):
                d = (float(Nbp)/n[i])*float(Lds)*(delta[i]/rref + 1)
                helix_stretch_energy += 0.5*S*n[i]*((d-Lds)**2)/Lds
                helix_bending_energy += 0.5*B*n[i]*Lds/((rref + delta[i])**2)
        helix_total_energy.append(helix_stretch_energy + helix_bending_energy)
bending_angle = (helix_total_energy.index(min(helix_total_energy))+1)
print("The bending angle of the bridge is",bending_angle, ", and the total energy is",\
              min(helix_total_energy)/(kb*T), "kBT.")
```

Calculations for open, tension-loaded clamps

With ssDNA strings hold the clamp open, the total energy stored in the tension-loaded clamps was calculated as the sum of energy of the bridge and the four ssDNA strings. For simplicity, we assume that the bridge remains an arc of circle in the open conformation.

The tension of the ssDNA strings was estimated based on the worm-like-chain (WLC) model[11] as

$$F_{WLC}(x) = \frac{k_b \cdot T}{L_{pss}} \cdot \left[ \frac{1}{4 \cdot \left(1 - \frac{x}{L_C}\right)^2} - \frac{1}{4} + \frac{x}{L_C} \right]$$

where $k_b$ is the Boltzmann constant, $T$ is the temperature (298 K), $L_{pss}$ is the persistence length of ssDNA, $L_C$ is the contour length of the ssDNA strings, and $x$ is the extension of the ssDNA strings.

The energy stored in the ssDNA strings was calculated by

$$E_{ssDNA\ strings}(\Delta x) = \sum_i \int_0^{\Delta x} F_{WLC}(x)\, dx$$

where the index $i$ iterates over all four ssDNA strings.

We calculated the total energy as a function of the bending angle ($\theta$) of the bridge. Using the geometrical relationship shown in **Box S1**, the end-to-end distance ($L_1$ and $L_2$) of the ssDNA strings can be calculated as

$$L_1 = 2 \cdot (R + d_1) \cdot \sin\frac{\theta}{2}$$

$$L_2 = 2 \cdot (R + d_2) \cdot \sin\frac{\theta}{2}$$

$$R = \frac{L}{\theta}$$

where $R$ denotes the radius of the bridge, $\theta$ denotes the bending angle of the bridge, $d_1$ or $d_2$ denotes the distance between the end of bridge and the end of the lower or upper ssDNA strings, and $L$ denotes the length of the bridge.

**Box S1**. Calculation of end-to-end distance of the ssDNA strings against the bending angle of the bridge.

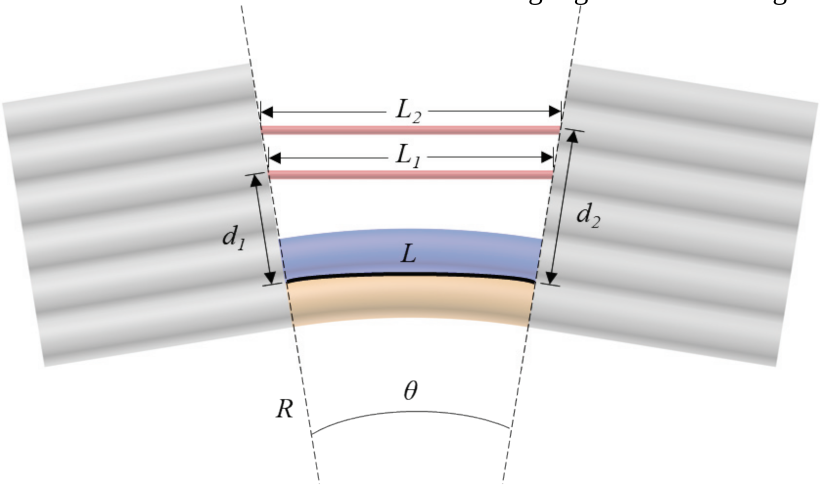


The total energy of the tension-loaded clamps can be calculated as

$$E_{total} = E_{bridge} + E_{ssDNA\ strings}$$

This calculation was done using the following Python (v 3.7.3 with SciPy[11]) script.

```
# Calculations for tension-loaded clamps

# The purpose of this script is to find the minimum energy of sum of bending
# energy of the bridge and stretching energy of the four ssDNA strings,
# and calculate the bending angle of the bridge, tension force in the ssDNA strings,
# end-to-end length of the ssDNA strings.


from numpy import sin, cos, pi, sqrt, inf
from scipy.integrate import quad
from math import ceil


# All constants are listed below

D = 2.5E-9 # Diameter of dsDNA (m)
Lss = 0.63E-9 # length of ssDNA (m/base)
Lds = 0.335E-9 # length of dsDNA (m/base)
kb = 1.38E-23 #Boltzmann constant (J/K)
T = 298 # Temperature(K)
Lpds = 50E-9 # persistence length of dsDNA (m)
Lpss = 0.75E-9 # persistence length of ssDNA (m)
S = 660E-12 # Stretch moduli of DNA helix (N)
B = 230E-30 # Bending moduli of DNA helix (N*m2)


# All adjustable parameters are listed below
```

```
N1 = 43 # number of nucleotides in the lower ssDNA strings (N1=43,N2=45 for when indelbp = 3 or 5)
N2 = 45 # number of nucleotides in the upper ssDNA strings (N1=34,N2=35 for when indelbp = 7 or 9 or
11)
e_sd1 = 6.25E-9 # distance between the end of the bridge and the end of the lower ssDNA strings (m)
e_sd2 = 8.75E-9 # distance between the end of the bridge and the end of the upper ssDNA strings (m)
Nss1 = 2 # number of the lower ssDNA strings
Nss2 = 2 # number of the upper ssDNA strings
Nds = 4 # number of helices in the bridge
Nbp = 42 # number of bp in the bridge (axial: no insertion or deletion)
L_beam = Nbp*Lds # length of the bridge
L_string1_max = N1*Lss # maximum length of the lower ssDNA string
L_string2_max = N2*Lss # maximum length of the upper ssDNA string
theta_upper_limit = L_beam/e_sd2 # maximum bending angle of the bridge
theta_step = 0.1 # step of bending angle of the bridge (in degree)
indelbp = 3 # bp(n) is the bp number of insertion or deletion from each outer or inner layer of
helices (Note: n can be 3, 5, 7, 9, or 11)
delta_coefficient = 1.25E-9 # dsDNA helix diameter coefficient

n = [0 for i in range(Nds)] # n is the number of basepairs installed in helices
n[0] = Nbp + indelbp
n[1] = Nbp + indelbp
n[2] = Nbp - indelbp
n[3] = Nbp - indelbp

delta = [0 for i in range(Nds)] # delta is the distance in nm from the center of mass midline
bending axis
delta[0]  = 1.0
delta[1]  = 1.0
delta[2]  = -1.0
delta[3]  = -1.0

for i in range(Nds):
          delta[i] *= delta_coefficient


# Worm-Like Chain Model predicts the force and energy stored in ssDNA as a function of end-to-end
distance of a ssDNA

def F(x,N):
    return((kb*T/Lpss)*(1/(4*((1-x/(N*Lss))**2))-0.25+x/(N*Lss)))


# Stretching energy stored in ssDNA strings

def Energy_string(x,N):
    return(quad(F,0,x,args=(N))[0])


# Calculate length of ssDNA string against bending angle theta, 1 for lower, 2 for upper

def L_string1(theta):
    return(2*(L_beam/theta+e_sd1)*sin(theta/2))

def L_string2(theta):
    return(2*(L_beam/theta+e_sd2)*sin(theta/2))


# Calculate the bending angle without ssDNA strings

helix_total_energy = []
for angle_in_degrees in range(1,180):
        angle_in_radians = angle_in_degrees/180.0*pi
        rref = L_beam/angle_in_radians
        helix_stretch_energy = 0
```

```
        helix_bending_energy = 0
        for i in range(Nds):
                d = (float(Nbp)/n[i])*float(Lds)*(delta[i]/rref + 1)
                helix_stretch_energy += 0.5*S*n[i]*((d-Lds)**2)/Lds
                helix_bending_energy += 0.5*B*n[i]*Lds/((rref + delta[i])**2)
        helix_total_energy.append(helix_stretch_energy + helix_bending_energy)
bending_angle = (helix_total_energy.index(min(helix_total_energy))+1)
print("The tension-free bending angle is",bending_angle)


# Calculate the total energy at different bending angles

if(N1*Lss <= L_beam):
        print("Please give a nt number in ssDNA string larger than: ",ceil(Nbp*Lds/Lss))
else:
        theta_all = []
        L_string1_all = []
        force1_all = []
        L_string2_all = []
        force2_all = []
        total_energy_all = []
        theta = theta_step # Starting from a very small theta, here choosing theta_step

        while(theta < bending_angle and L_string1(theta/180.0*pi)<L_string1_max\
              and L_string2(theta/180.0*pi)<L_string2_max):
                rref = L_beam/(theta/180.0*pi)
                beam_stretch_energy = 0
                beam_bending_energy = 0
                for i in range(Nds):
                        d = (float(Nbp)/n[i])*float(Lds)*(delta[i]/rref + 1)
                        beam_stretch_energy += 0.5*S*n[i]*((d-Lds)**2)/Lds
                        beam_bending_energy += 0.5*B*n[i]*Lds/((rref + delta[i])**2)
                total_energy = beam_stretch_energy + beam_bending_energy\
                               + Nss1*Energy_string(L_string1(theta/180.0*pi),N1)\
                               + Nss2*Energy_string(L_string2(theta/180.0*pi),N2)
                theta_all.append(theta)
                L_string1_all.append(L_string1(theta/180.0*pi))
                force1_all.append(F(L_string1(theta/180.0*pi),N1))
                L_string2_all.append(L_string2(theta/180.0*pi))
                force2_all.append(F(L_string2(theta/180.0*pi),N2))
                total_energy_all.append(total_energy)
                theta += theta_step
        index = total_energy_all.index(min(total_energy_all))
        print("The minimum energy point has the bending angle of", theta_all[index],\
              "degree, the lower ssDNA string length of", L_string1_all[index]*1.0E9,\
              "nm, with", N1, "nt in the string, the force in lower ssDNA strings of",\
              force1_all[index]*1.0E12, "pN; the upper ssDNA string length of", \
              L_string2_all[index]*1.0E9, "nm, with", N2,\
              "nt in the string, the force in upper ssDNA strings of",\
              force2_all[index]*1.0E12, "pN, and the total energy of",\
              total_energy_all[index]/(kb*T), "kBT.")
```

We assume the clamps tubulate vesicles by releasing energy stored in the bridge during the open-to-closed reconfiguration. Therefore, it is useful to calculate the energy difference of bridge between the closed and open conformations, as follows:

$$\Delta E = E_{bridge\_closed} - E_{bridge\_open}$$

This calculation was done using the following Python (v 3.7.3 with SciPy[11]) script.

```
# Calculations for energy released from the open-to-closed reconfiguration

from numpy import pi, sqrt
```

```
# All constants are listed below

D = 2.25E-9 # Diameter of dsDNA (m)
Lds = 0.335E-9 # length of dsDNA (m/base)
kb = 1.38E-23 #Boltzmann constant (J/K)
T = 298 # Temperature(K)
S = 660E-12 # Stretch moduli of DNA helix (N)
B = 230E-30 # Bending moduli of DNA helix (N*m2)


# All adjustable parameters are listed below

Nds = 4 # number of helices in the bridge
Nbp = 42 # number of bp in the bridge without insertion or deletion
L_conn = Nbp*Lds # length of the bridge
indelbp = 3 # bp(n) is the number of bp inserted or deleted from each outer or inner layer of
helices (Note: n can be 3, 5, 7, 9, or 11)
degree_open = 3 # bending angle of open, tension-loaded clamp
degree_closed = 38 # bending angle of closed, tension-released clamp
delta_coefficient = 1.25E-9 # each DNA helix is modeled as a 2.5 nm wide rod

n = [0 for i in range(Nds)] # n is the number of bp installed in helices
n[0] = Nbp + indelbp
n[1] = Nbp + indelbp
n[2] = Nbp - indelbp
n[3] = Nbp - indelbp

delta = [0 for i in range(Nds)] # distance of a helix from the bending axis (nm)
delta[0]  = 1.0
delta[1]  = 1.0
delta[2]  = -1.0
delta[3]  = -1.0

for i in range(Nds):
          delta[i] *= delta_coefficient


# Calculate the energy released from the open-to-closed reconfiguration

helix_total_energy = []
for angle_in_degrees in range(1,180):
        angle_in_radians = angle_in_degrees/180.0*pi
        rref = L_conn /angle_in_radians
        helix_stretch_energy = 0
        helix_bending_energy = 0
        for i in range(Nds):
                d = (float(Nbp)/n[i])*float(Lds)*(delta[i]/rref + 1)
                helix_stretch_energy += 0.5*S*n[i]*((d-Lds)**2)/Lds
                helix_bending_energy += 0.5*B*n[i]*Lds/((rref + delta[i])**2)
        helix_total_energy.append(helix_stretch_energy + helix_bending_energy)
bending_angle = (helix_total_energy.index(min(helix_total_energy))+1)
print("The energy released from open-to-closed reconfiguration is", \
      (helix_total_energy[degree_closed-1]-helix_total_energy[degree_open-1])/(kb*T), "kBT.")
```

The calculation results for all five versions of clamps are summarized in **Box S2**.

**Box S2**. Summary of parameters of the DNA clamps.

| DNA clamps | indel (bp) | bending angle: closed* | bending angle: open* | No. of dTs in each string lower \| upper | Tension (pN) in each string lower \| upper | $\Delta E_{bridge}$ ($k_BT$)** |
|---|---|---|---|---|---|---|
| I | 3 | 36° (38°) | 17° (3°) | 43 \| 45 | 7.7 \| 7.1 | -6 |
| II | 5 | 62° (63°) | 22° (18°) | 43 \| 45 | 10.0 \| 10.1 | -22 |
| III | 7 | 85° (88°) | 20° (9°) | 34 \| 35 | 17.6 \| 17.3 | -63 |
| IV | 9 | 100° (113°) | 30° (15°) | 34 \| 35 | 21.7 \| 23.1 | -94 |
| V | 11 | 116° (137°) | 33° (21°) | 34 \| 35 | 25.9 \| 30.1 | -153 |

* Measured bending angle (theoretical bending angle).

** Maximum energy released by the bridge of DNA clamps from reconfiguration using measured bending angles (see above).

**Proposed mechanism of membrane tubulation**

Membrane tubulation is energetically costly at the two main steps: nucleation and elongation (**Box S3**)[12].

**Box S3**. Procedure of membrane tubulation. A tube with a radius of $r_t$ is pulled from a vesicle with a radius of $r_v$ to an extension of $l_t$. Tubulation is a 2-step process with a nucleation step (black dashed line) and an elongation step. The orange dashed line represents the 'foot' of the tube.

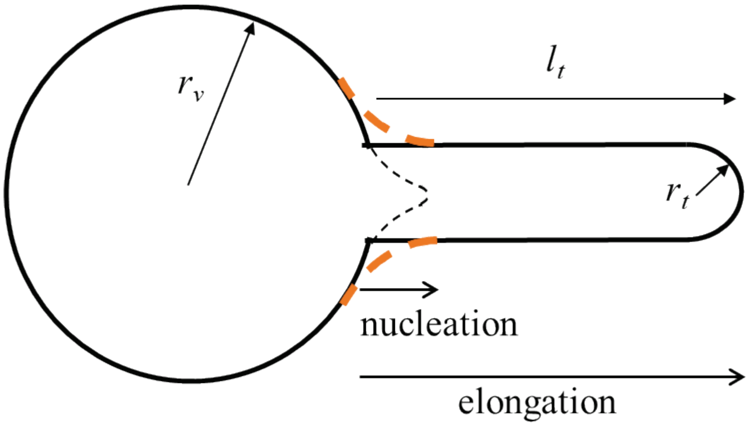


Nucleation of the tube is achieved by a pulling force leading to sufficient membrane deformation. It implies the formation of a small membrane spot, typically tens of nanometers, with a sharp curvature preventing the presence of several clamps on the nucleation spot. Hence, it is likely that a single clamp is responsible for the tube nucleation. The curvature energy cost for this tube budding is in the 25 – 30 $k_BT$ range (See "Nucleation of a tube" below for details). Comparing this value with the energy gain of each clamp type for the triggered open-to-close reconfiguration suggests that clamp I cannot provide enough energy while the other four types of clamps can (**Box S2**). This simple model provides an explanation for the low tubulation efficiency observed for clamp I. However, it cannot explain the low tubulation efficiencies for clamp IV and V with GUVs and all other clamps with LUVs.

These low tubulation rates can be predicted by considering the elongation stage. Tube elongation starts when the bud is long enough, a few nanometers to a micrometer depending on the surface tension (See "Nucleation of a tube" below for details). The energy involved in growing a unit of tube depends on the type of vesicles. For GUVs, the tube area is much smaller than the rest of the vesicle, making the energy cost for tube elongation almost only dependent on the tube bending energy, *i.e.* on the tube diameter. The corresponding energy values per unit area for tubes induced by each clamp are indicated in **Box S4**. The total energy provided by the clamps can be obtained from the released energy via open-to-close reconfiguration (**Box S2**), the clamp density (roughly estimated to be 610 clamps per µm² based on fluorescence microscopy data) and the fraction of correct (bent) clamps (**Figure S18**). The energy ratio shows that clamps IV and V do not provide enough energy for tube elongation. Hence, for GUVs, according to this model, only clamps II and III are able to nucleate and elongate tubes by triggering their conformational change. This prediction is consistent with the observations. For LUVs, the tube area can become commensurate with the vesicle area, making the tube elongation process more complex: as the tube grows, the surface tension of the membrane increases making tubulation more difficult to achieve (See "Elongation of a tube" for quantitative details). Considering the energy gain from the clamps and the clamp density, we can predict that, for 200-nm LUVs, the surface tension increase prevents the budding tube from growing further than a few dozens of nanometers for all clamps. Hence, this model predicts that tubes can hardly form in the case of LUVs, which also conforms with the observed low tubulation efficiency.

**Box S4**. Summary of the released energy provided by the clamp reconfiguration and the required tube bending energy associated with the corresponding tube radius. The ratio between the released energy and the tube bending energy per effective clamp area is present in the last column. A value larger than 1 indicates that the tube can elongate upon clamp reconfiguration. A value smaller than 1 suggests that tubulation is unlikely (clamps IV and V).

| clamp | released energy ($k_BT$)[a] | F fraction of correct (bent) clamps [b] | mean released energy ($k_BT$)[c] | $r_t$ (nm)[d] | tube bending energy ($k_BT$ per clamp molecular area)[e] | energy ratio[f] |
|---|---|---|---|---|---|---|
| I | -6 | 0.95 | -5.7 | 54 | 2 | 2.8 |
| II | -22 | 0.88 | -19.4 | 25 | 12 | 1.6 |
| III | -63 | 0.62 | -39.1 | 24 | 13 | 3.0 |
| IV | -94 | 0.31 | -29.1 | 14 | 50 | 0.58 |
| V | -153 | 0.30 | -45.9 | 13 | 58 | 0.79 |

[a] Released energy is maximum energy released by the bridge of DNA clamps from reconfiguration using measured bending angles (Box S2).
[b] F, or fraction of correct (bent) clamps, is obtained from **Figure S18**.
[c] Mean released energy is calculated by released energy times F.
[d] $r_t$ denotes the measured radius of tube.
[e] Tube bending energy denotes the required energy for tube elongation associated with the corresponding tube radius.
[f] Energy ratio is calculated by dividing the absolute value of mean released energy by tube bending energy.

The case of the closed, inherently tension-free (ITF) clamps on GUVs is more difficult to quantitatively model. Unlike open-to-close reconfiguration, tubulation can only come from the initial binding of the cholesterols for the ITF clamps. Each cholesterol insertion brings 25 $k_BT$ energy[13]. Hence, in theory, eight cholesterol moieties provide more than enough energy to trigger tubulation. However, this energy is not used in an effort to bend the membrane and most of it will be dissipated. In the case of the clamps with the largest curvatures, some cholesterol moieties may not even insert in the membrane. In addition, binding of the clamps is progressive, bringing the energy sequentially instead of simultaneously in the case of the triggered conformational change and preventing any cooperative effect on the elongation of the tube. This difficulty to transfer the cholesterol insertion energy to bending energy and the lack of cooperativity explain the low efficiency observed with ITF clamps.

Energetics of tube formation

In this section, we will quantitatively explain the forces and energies involved in the formation of a tube from a vesicle with a surface tension $\sigma$ and a bending modulus $k_c$ when a local pulling force $f$ is applied to the vesicle membrane. Following the notations presented in reference[12], we normalize all lengths by

$R_0 = \sqrt{\frac{k_c}{2\sigma}}$ and forces by $f_0 = 2\pi\sqrt{2\sigma k_c}$. A typical value for $k_c$ is 10 $k_B T$ and, to pull a tube, $\sigma$ is between $10^{-7}$ and $10^{-4}$ N/m. Hence $R_0$ (membrane tube radius at equilibrium) ranges between 15 and 150 nm and $f_0$ (pulling force at equilibrium) is between 0.2 and 20 pN.

Nucleation of a tube

To form a tube, the force $f$ must be large enough to generate a sufficient deformation to overcome the energy barrier for nucleation[12]. This force, $f_{max}$, is about 1.13 $f_0$ and the local deformation, $l(d)$, of the (flat) vesicle membrane at a distance $d$ from the pulling spot can be written:

$$\frac{l(d)}{R_0} = 2\frac{f_{max}}{f_0}\left[ln\left(\frac{d}{\sqrt{2}R_0}\right) + K_0\left(\frac{d}{\sqrt{2}R_0}\right)\right]$$

where $K_0(x)$ is the modified Bessel function of the second kind. The maximum length of this nucleating tube, $l_{max} = l(0)$, is about 8 $R_0$, *i.e.* up to 1 µm.

The curvature energy associated with the shape of the membrane upon tube nucleation can be readily integrated and is about 2.8 $k_c$, *i.e.* in the range 25 – 30 $k_B T$ for usual $k_c$ values. All clamps except clamp I are able to provide this energy via open-to-close reconfiguration (**Box S2**).

Elongation of a tube

As the tube is being elongated after nucleation, the constant lumen volume force imposes that the radius of the vesicle $r_v$ decreases with the tube length, $l_t$, increase. This decrease depends on the tube radius $r_t$ and can be expressed as:

$$r_v(l_t) = \left(r_{v_0}^3 - \frac{3}{4}r_t^2\left(l_t - l_{t_0}\right)\right)^{1/3}$$

where the subscript '0' denotes a reference elongation.

Ignoring the contribution of the foot of the tube (orange dashed line in **Box S3**), the free energy $F$ can be written:

$$F(l_t) = \sigma A + \iint_S \frac{k_c}{2}(2\langle c\rangle)^2 dS$$

In addition, $\sigma = \frac{\kappa(A-A_0)}{A} + \sigma_0$, where $\kappa$ is the elastic modulus, and the mean curvature $\langle c\rangle$ is $\frac{1}{r_v}$ on the vesicle and $\frac{1}{2r_t}$ on the tube.

Hence, the free energy variation from the reference elongation is:

$$\Delta F(l_t) = \pi\left[4\kappa\left({r_v}^2 - r_{v_0}^2\right) + 2\kappa r_t\left(l_t - l_{t_0}\right) + k_c\left(2\frac{r_t^2}{r_{v_0}^2} - 2\frac{r_t^2}{{r_v}^2} + \frac{\left(l_t - l_{t_0}\right)}{r_t}\right)\right]$$

For a GUV, the vesicle area is much larger than that of the tube. In this case, the last term of the free energy variation dominates and $\Delta F$ can be written:

$$\Delta F(l_t) \approx \pi k_c \frac{\left(l_t - l_{t_0}\right)}{r_t}$$

This energy needs to be compared to the energy provided by the clamps. Experimentally, we measured $c \approx$ 610 clamps per µm². Only a fraction of these clamps, $F$, are correctly closed during open-to-close reconfiguration (**Box S4**). Hence, the effective clamp molecular area is $c/F$, which corresponds to a tube length change, $l_t - l_{t0} = \frac{c}{2\pi F r_t}$, and a free energy:

$$\Delta F(l_t) \approx k_c \frac{c}{2F{r_t}^2}$$

This free energy, indicated in **Box S4**, cannot be provided by clamps IV and V.

For LUVs, the tube area represents a significant fraction of the total area. Hence, the surface tension term in equation

$$\Delta F(l_t) = \pi \left[ 4\kappa\left({r_v}^2 - r_{v0}^2\right) + 2\kappa r_t\left(l_t - l_{t0}\right) + k_c\left(2\frac{r_t^2}{r_{v0}^2} - 2\frac{r_t^2}{{r_v}^2} + \frac{\left(l_t - l_{t0}\right)}{r_t}\right)\right]$$

needs to be kept. The resulting surface tension is presented in **Box S5**. The results show that forming a tube from LUVs with the clamps is impossible without reaching the lysis tension within a few dozens of nanometers unless the LUV is initially very deflated, *i.e.* there is enough excess of surface for surface tension to remain null during the tubulation process.

**Box S5**. Surface tension increase associated with a change in tube length, $l_t - l_{t_0}$, when the vesicle is initially spherical without any significant surface tension for a tube with a 20 nm radius, a 10 $k_BT$ bending modulus, a 50 mN/m elastic modulus. The case of a 200-nm LUV (blue) and a 10-µm GUV (orange) is presented. The lysis tension is typically 10 mN/m. This curve indicates that a tube cannot significantly extend from an LUV.

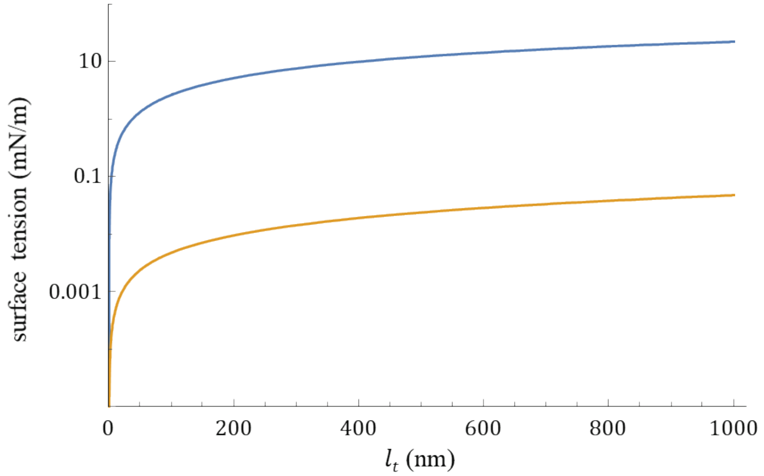


Tension-driven tube diameter

For GUVs, the tube radii $r_t$, closely match the tension-released clamp radii, $r_c$ (**Figure 4d**), indicating the clamps are fully controlling the tube geometry. In the case of LUVs, the tube radii are always in the range 15-20 nm, showing that even though the clamps drive tubulation, another parameter controls the tube dimensions. In the section “elongation of a tube” above, we saw that the fast increase of surface tension limits the length of the tube formed from LUVs. It is well-established that the tube radius decreases with the surface tension according to the following equation[12], hinting that surface tension may also be responsible for the observed small tube radii in LUVs:

$$r_t = \sqrt{\frac{k_c}{2\sigma}}$$

The predicted tube radius (**Box S6**) suggests that the surface tension after tubulation from LUVs is above $10^{-4}$ N/m.

**Box S6**. Predicted tube radius with the vesicle surface tension. The bending modulus was chosen at 10 $k_BT$.

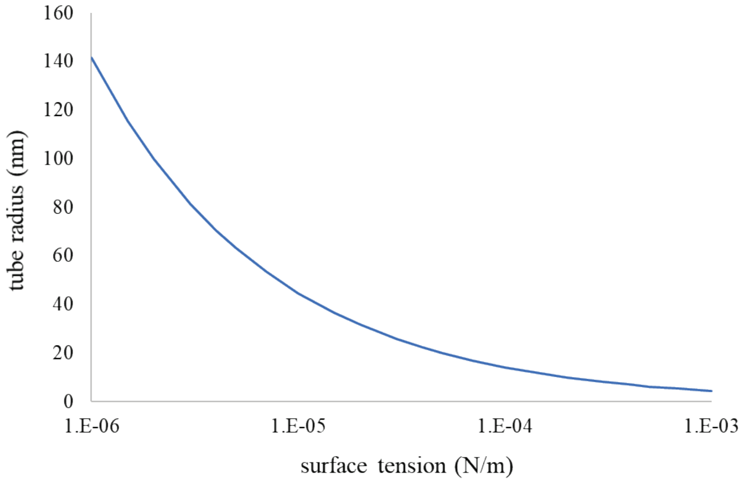

#### SUPPLEMENTAL FIGURES

**Figure S1.** caDNAno design diagrams of an open, tension-loaded DNA clamp (a) and a closed, inherently tension-free DNA clamp (b). Blue: scaffold strand; grey: core staple strands; green: handles for cholesterol modification. Purple: handles for Alexa Fluor 647 labeling. Red: ssDNA strings. The staples in the black box were added in step 2 of the 2-step assembly (see "Design and assembly of DNA-origami structures").

(a)

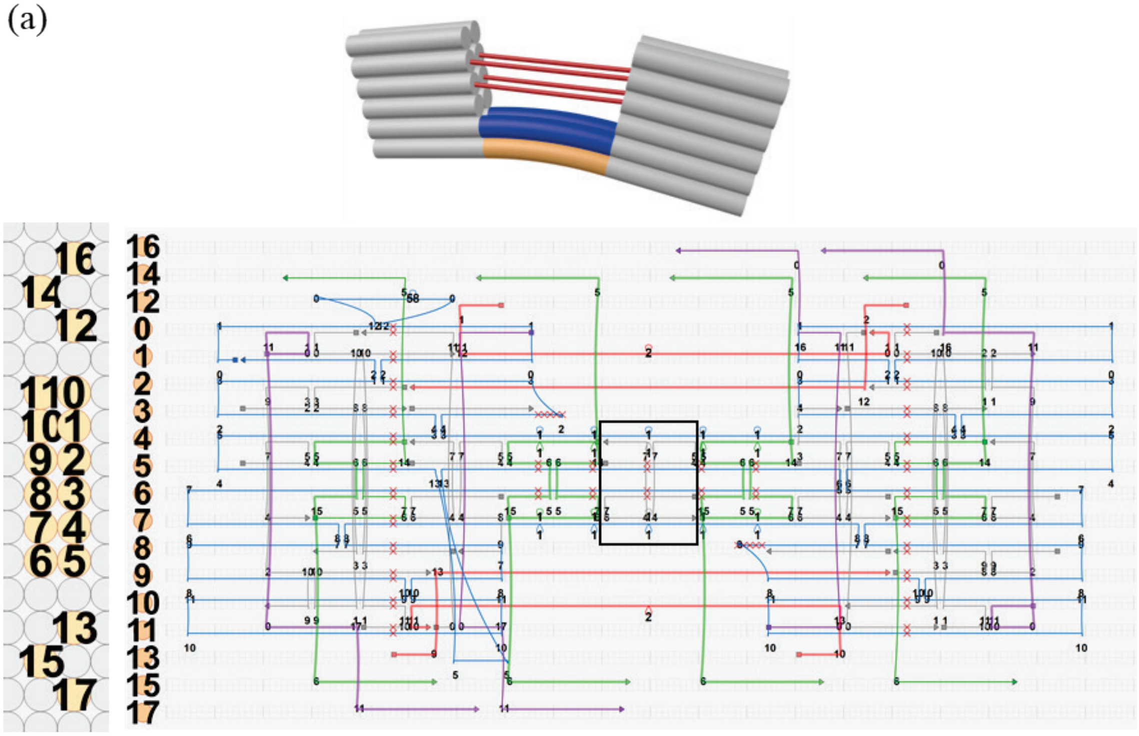

(b)

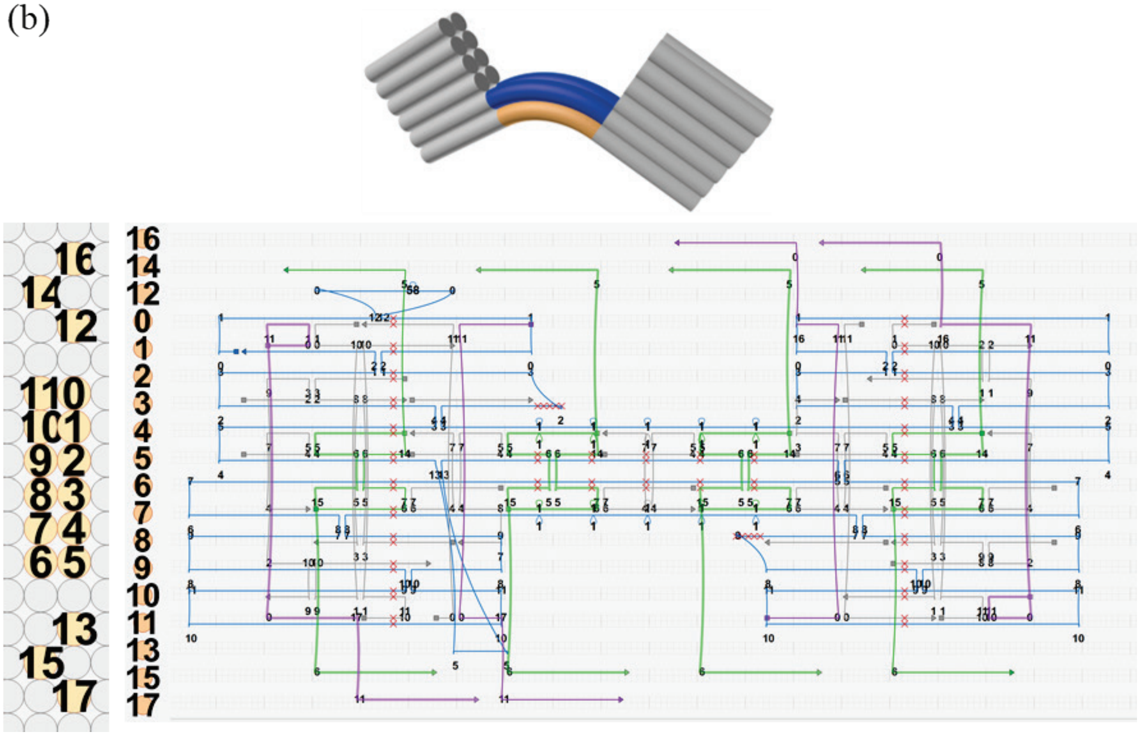

**Figure S2.** Agarose gel (1.5%) electrophoresis of DNA clamps assembled in one pot at various $Mg^{2+}$ concentrations. Gel and TEM images indicate that tension-loaded DNA clamps tend to form dimers (left), whereas the inherently tension-free clamps mostly fold as monomers (right). Scale bars: 50 nm.

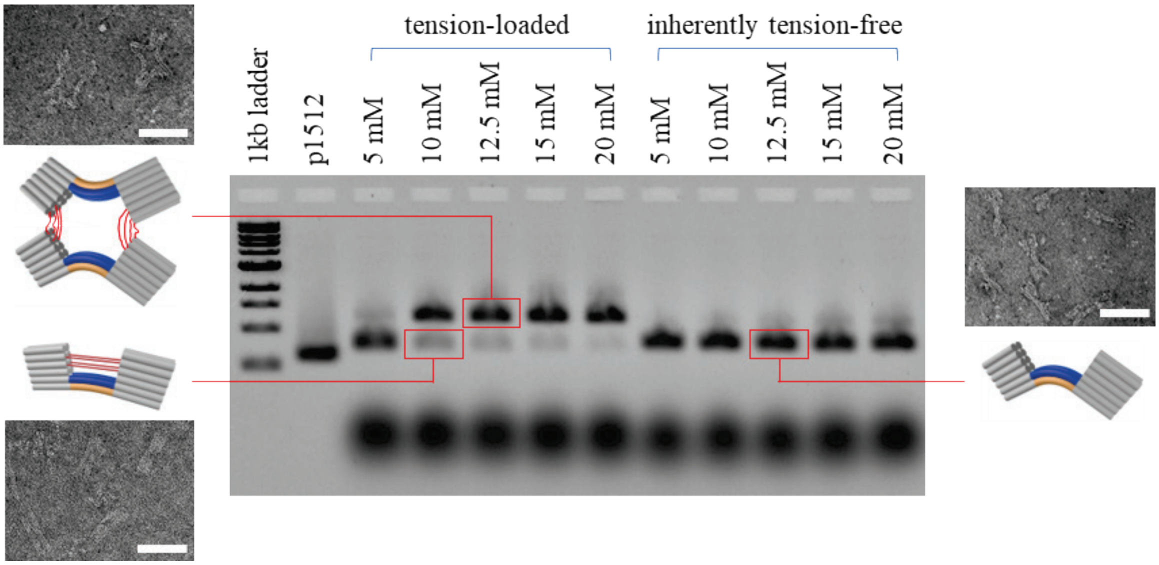

**Figure S3.** Agarose gel (2%) analysis of tension-loaded DNA clamps assembled in two steps. “Purified” lane contains two-step assembled DNA clamps after purification by PEG fractionation. Scale bar: 50 nm.

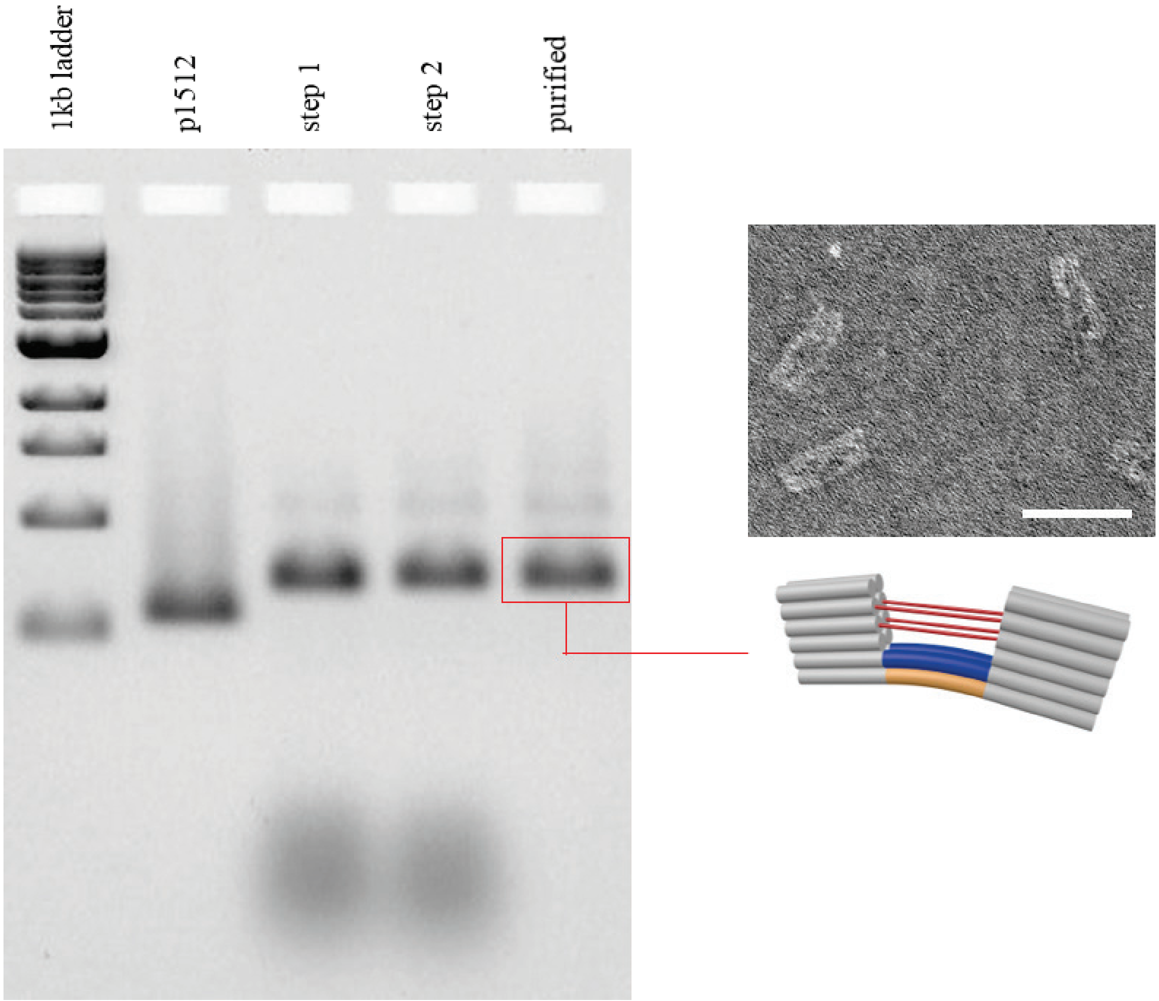

**Figure S4.** caDNAno design diagrams of DNA extenders. Blue: scaffold strand; grey: core staple strands; black: staple strands at the end of the structure for enabling or blocking end-to-end joining.

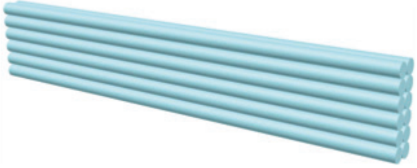

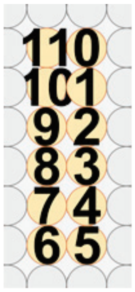


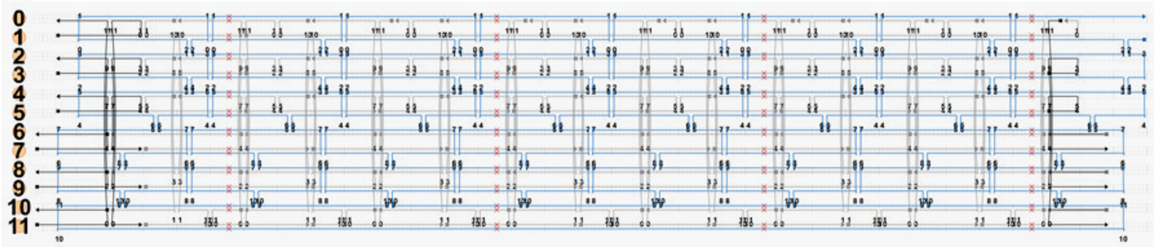

left extender

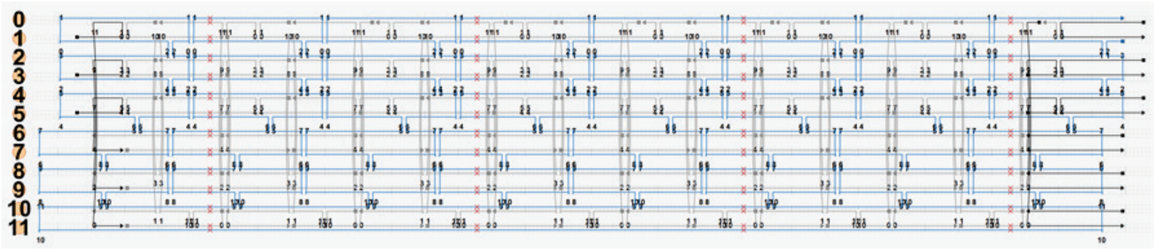

right extender

**Figure S5.** TEM images of DNA extenders after purification by PEG fractionation. Scale bars: 50 nm.

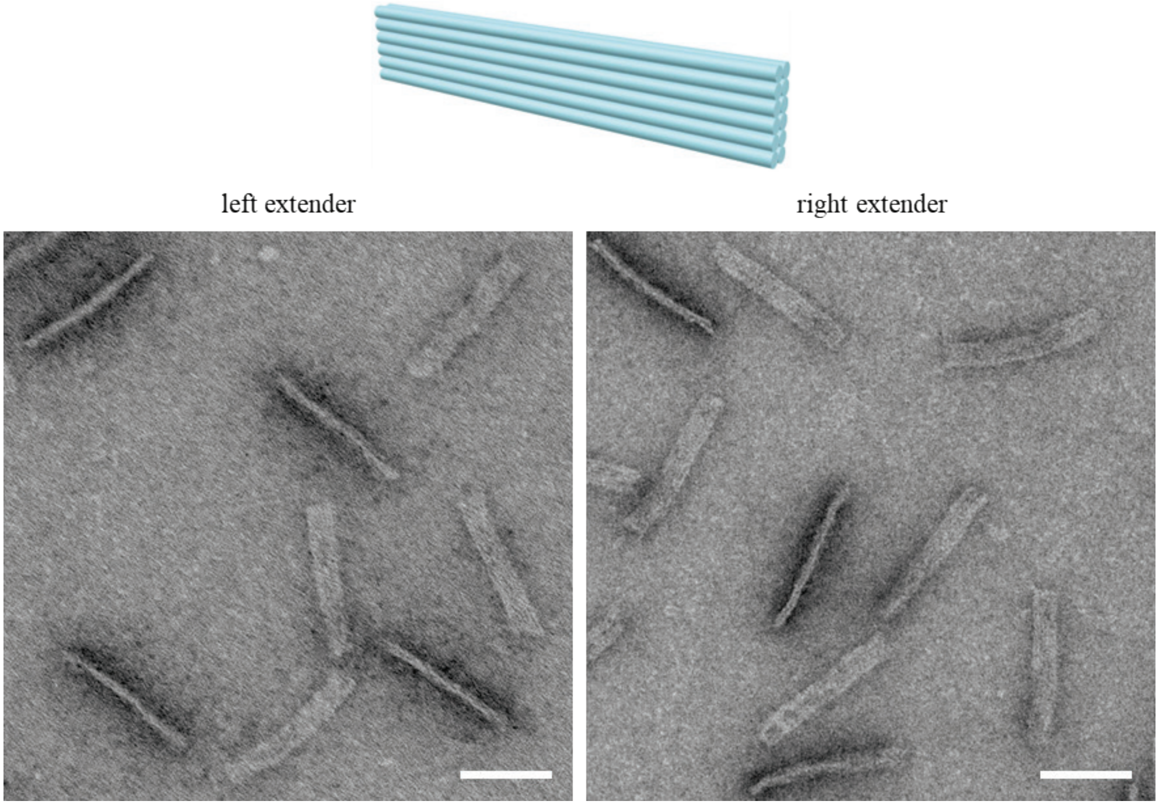

**Figure S6.** Binding affinity of open, tension-loaded DNA clamps towards LUVs. (a) Results of an isopycnic centrifugation (float up assay) of a mixture of DNA clamps and LUVs. Left: A schematic of the iodixanol gradient, with the bottom layer (yellow) containing DNA clamps and LUVs, formed before centrifugation. Right: fractions 1–16 collected from the top to the bottom of the gradient solution after centrifugation imaged in the rhodamine and Alexa Fluor 647 channels. (b) Agarose gel (1.5%, containing 0.05% SDS) electrophoresis study of all 16 fractions shown in (a). Co-existence of DNA and lipid in fractions 1–8 (cyan) confirmed binding between DNA clamps and vesicles. Pseudo-color fluorescence: green = rhodamine (Rhod), red = Alexa Fluor 647 (Alexa647).

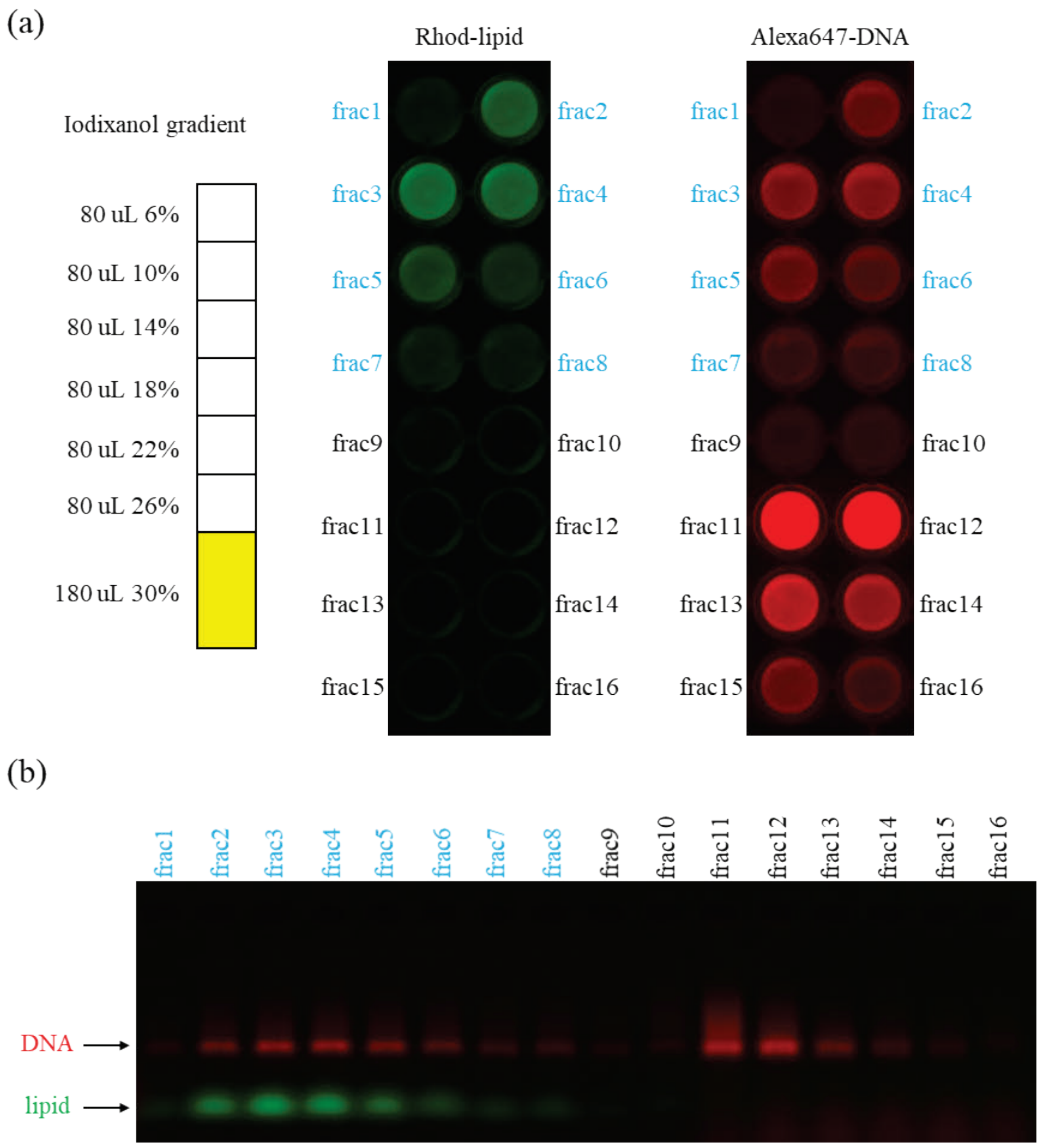

**Figure S7.** Tubulation efficiency of LUVs as a function of theoretical membrane coverage. N.D.: not detected.

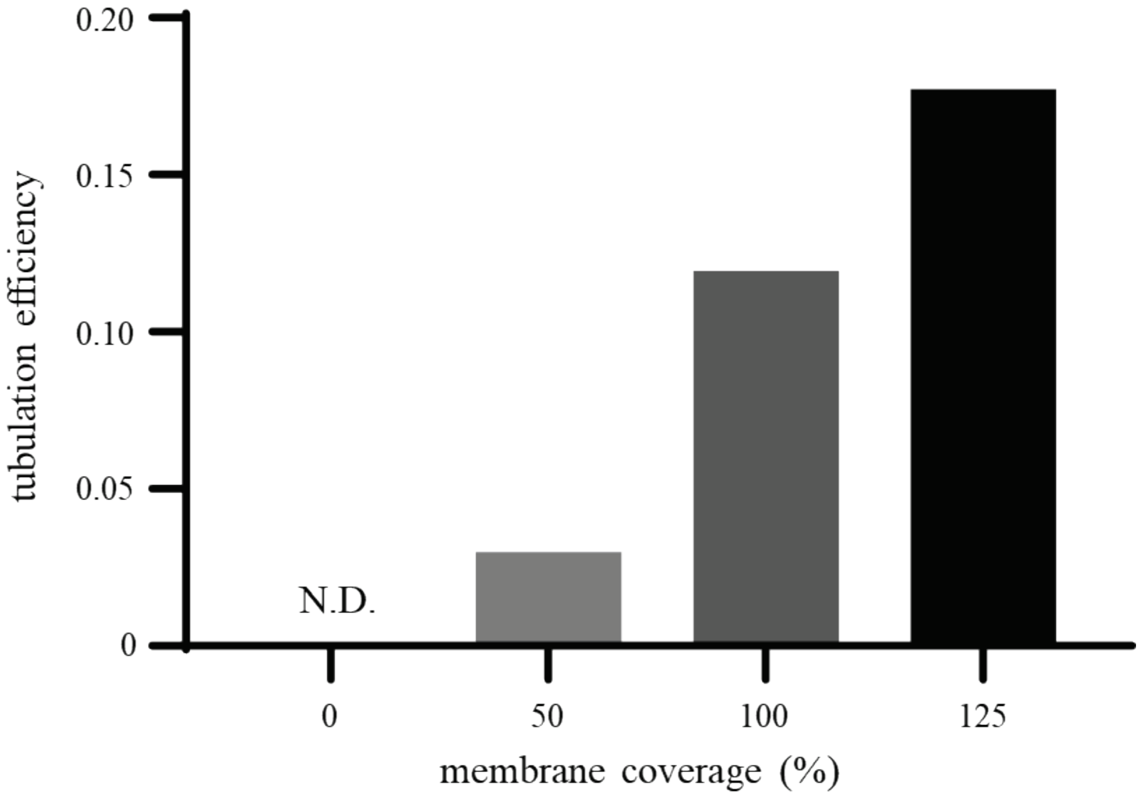

**Figure S8.** Time-course study of two incidents (a and b) of membrane tubule breaking after DNase I treatment. Scale bars in confocal microscopy images: 10 μm.

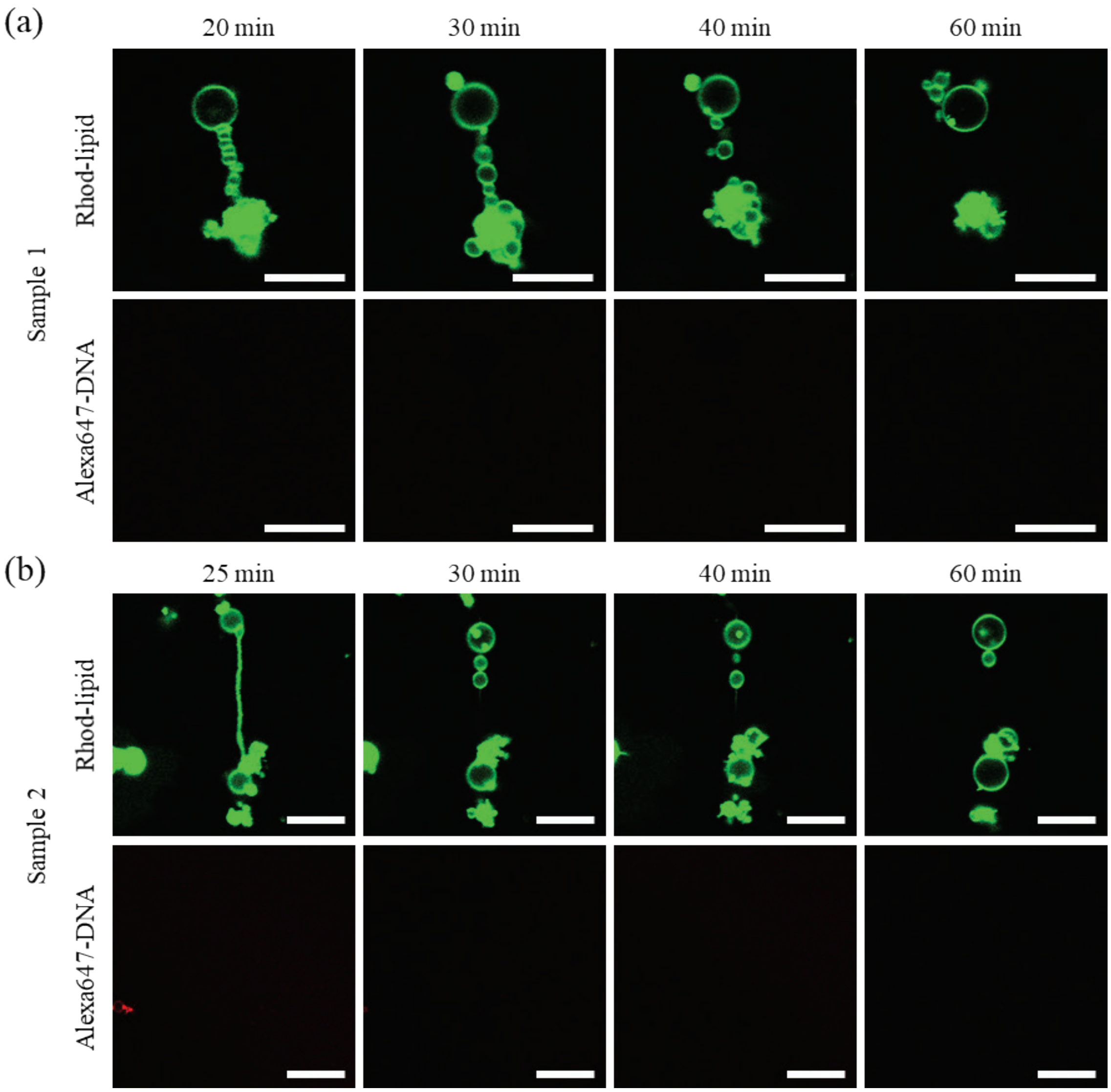

**Figure S9.** Time-course study of DNase I digestion of DNA clamps in solution. (a) Agarose gel (1.5%) electrophoresis of DNA clamps co-incubation with DNAse I for 5 min – 1 hour. Starting from left, lane 1: 1 kb DNA ladder; lane 2: Alexa Fluor 647-labeled DNA clamps without cholesterol (chol) modification ran as a dense band; lane 3: Cholesterol-modified, Alexa Fluor 647-labeled DNA clamps ran tardily and as a smear; lanes 4–10: Cholesterol-modified, Alexa Fluor 647-labeled DNA clamps after incubation with DNase I for different amount of time. Pseudo-color fluorescence: green = EtBr, red = Alexa Fluor 647. (b) EtBr fluorescence intensity of DNA clamps after DNase I treatment (Lane 4–10 in a) normalized to the untreated clamps (Lane 3 in a).

(a)

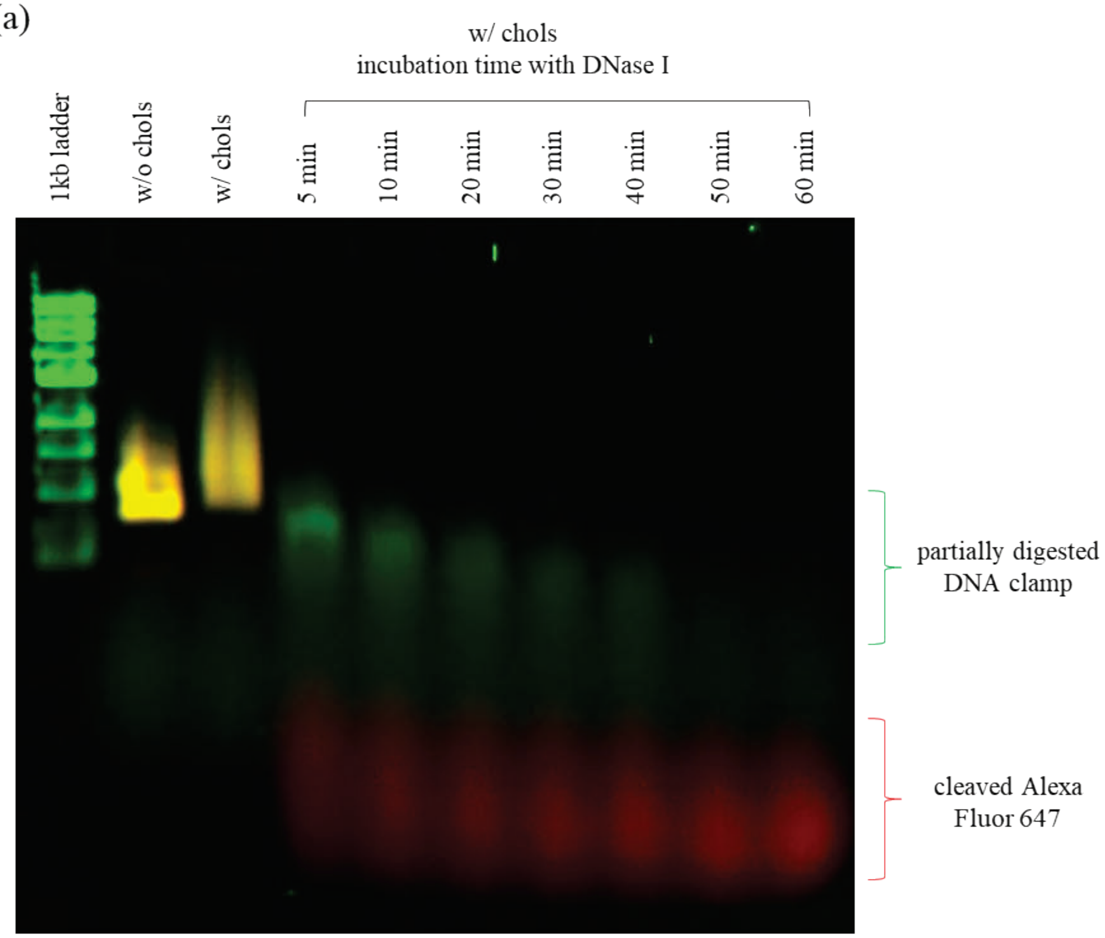


(b)

| incubation time (min) | 0 | 5 | 10 | 20 | 30 | 40 | 50 | 60 |
|---|---|---|---|---|---|---|---|---|
| normalized intensity | 100.0% | 39.4% | 30.4% | 19.8% | 11.7% | 7.2% | 1.0% | 0.5% |

**Figure S10.** The bridge designs of DNA clamps I–V.

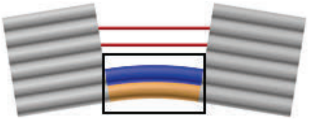

| DNA clamps | bending angle | # of indel bps | bp indel pattern |
|---|---|---|---|
| I | 38° | 3 | 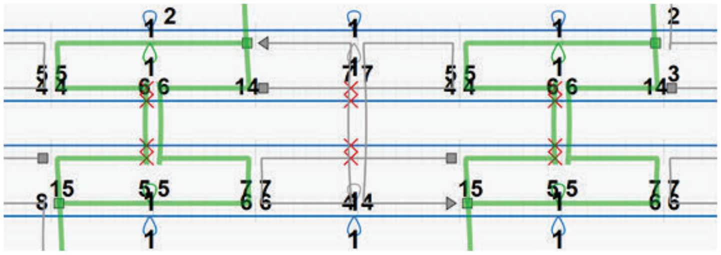 |
| II | 63° | 5 | 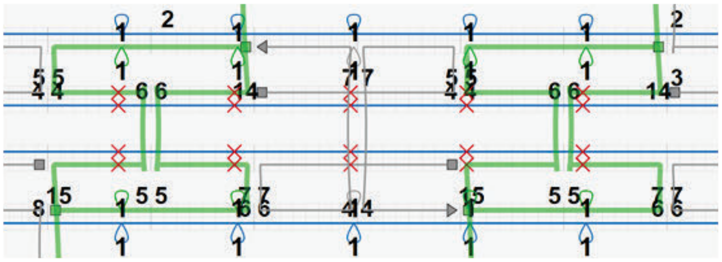 |
| III | 88° | 7 | 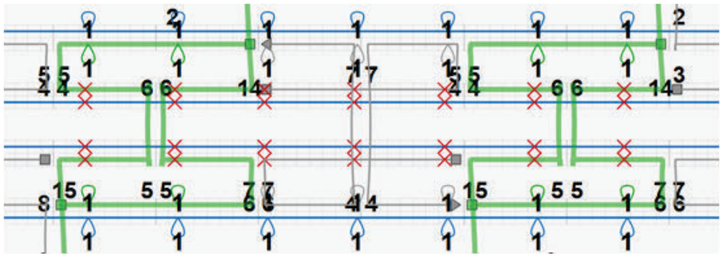 |
| IV | 113° | 9 | 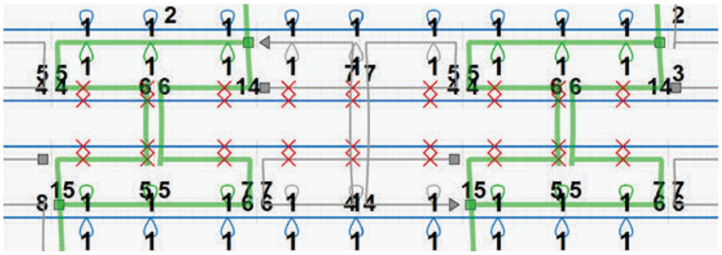 |
| V | 137° | 11 | 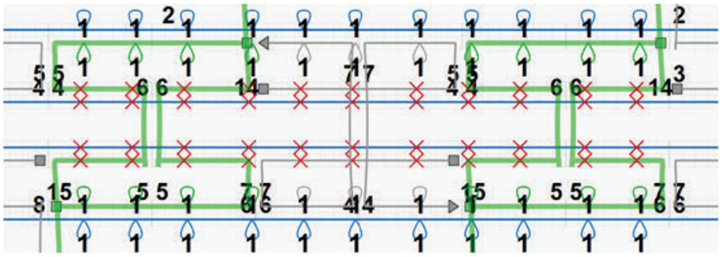 |

**Figure S11.** Revised staple strand design in the bridge for improved folding efficiency. (a) caDNAno design diagrams of the staple strands (grey) added at the second step of the 2-step folding. Left: original design (step 2), where each staples traverses three helices; right: revised design (step 2 rev), where each staple traverses only two helices, binding more stably (>12 bp) in each helix. (b) Agarose gel (2%) electrophoresis study of tension-loaded DNA clamps. Note the reduction of multimers in clamps VI and V (highlighted in red boxes) when folding with using "step 2 rev" staple strands.

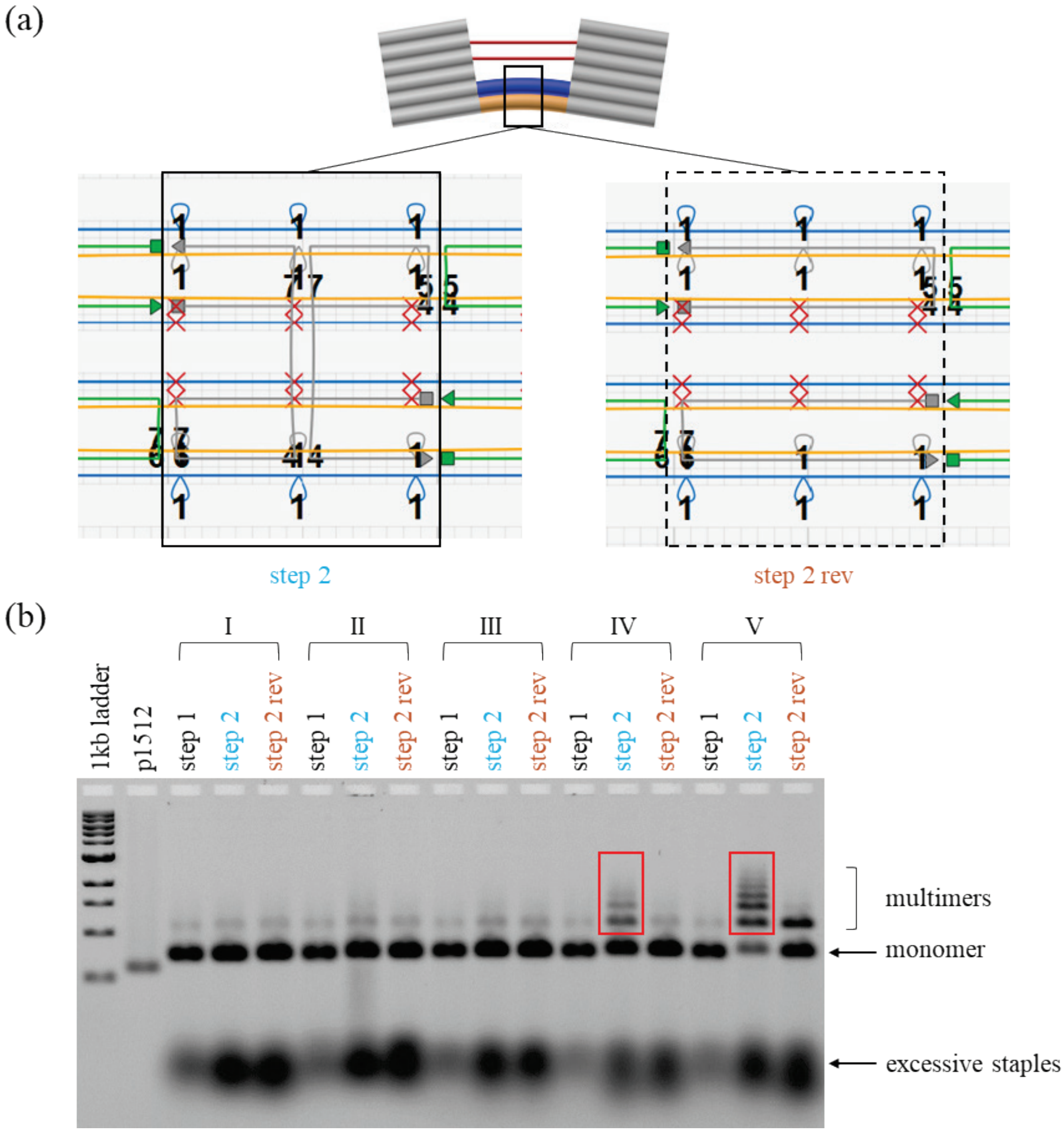

**Figure S12.** Class averaging of selected particles for DNA clamp I. (a) Open (tension-loaded) state, (b) closed (tension-released) state, (c) closed (inherently tension-free) state. Box size: 93.9 nm.

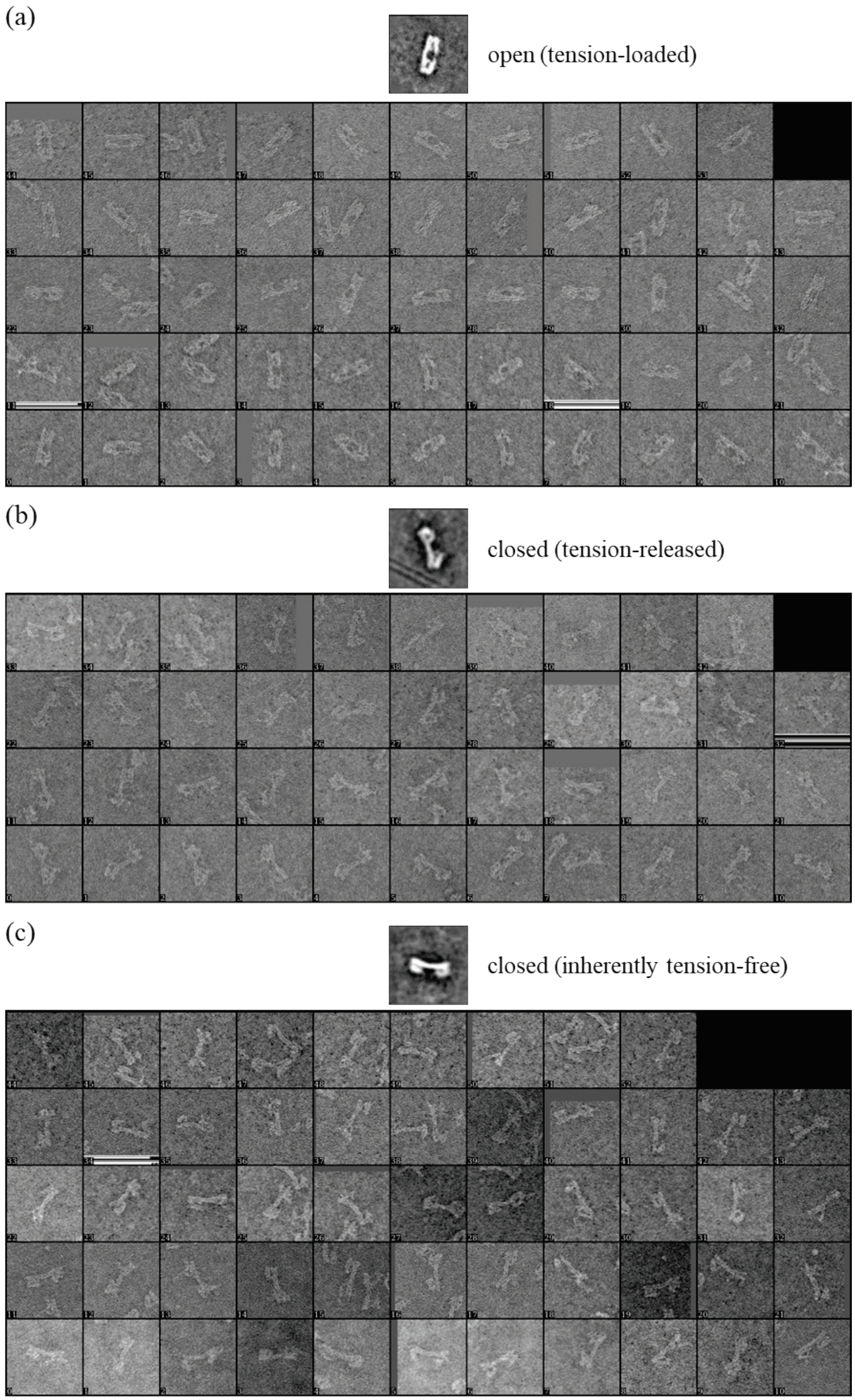

**Figure S13.** Class averaging of selected particles for DNA clamp II. (a) open (tension-loaded) state. (b) closed (tension-released) state. (c) closed (inherently tension-free) state. Box size: 93.9 nm.

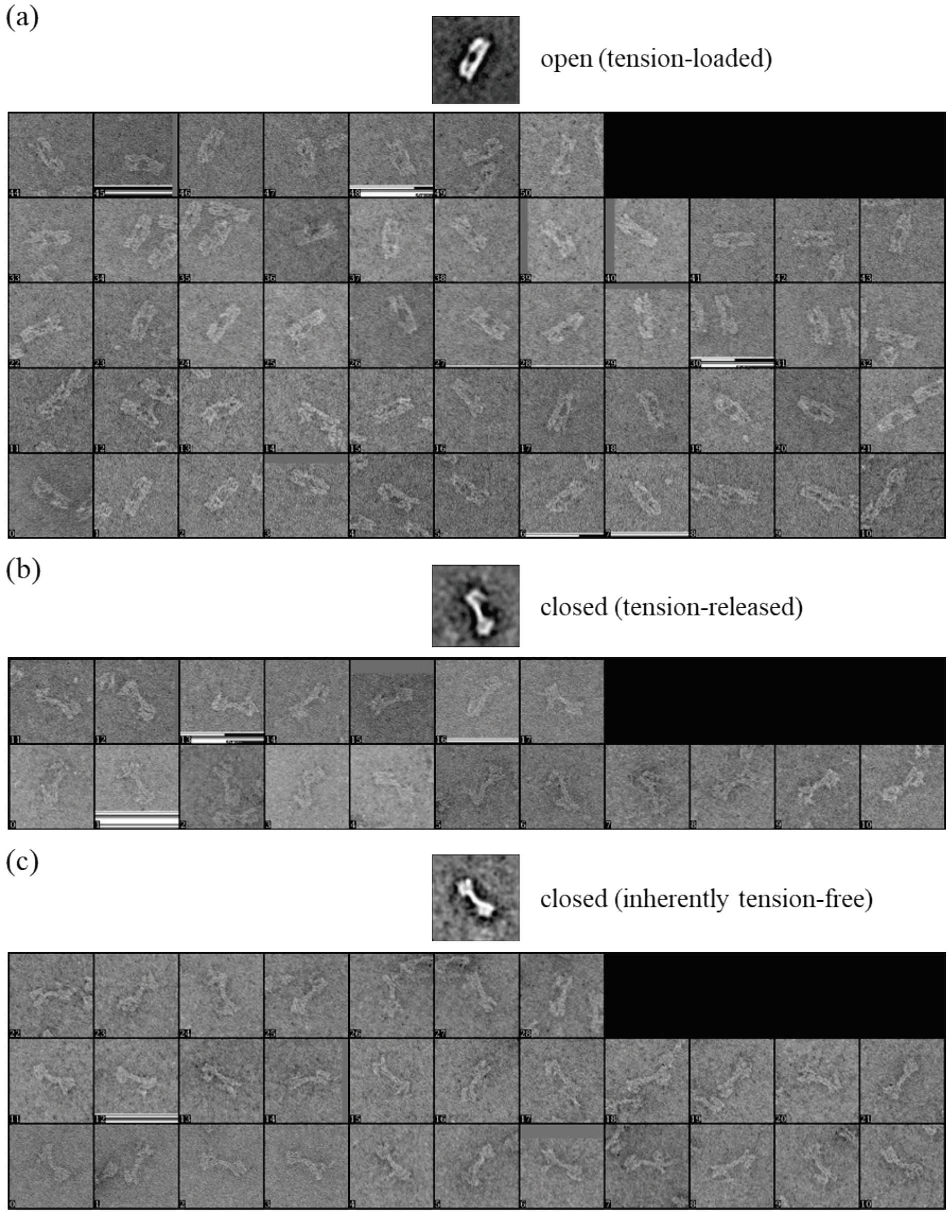

**Figure S14.** Class averaging of selected particles for DNA clamp III. (a) open (tension-loaded) state. (b) closed (tension-released) state. (c) closed (inherently tension-free) state. Box size: 93.9 nm.

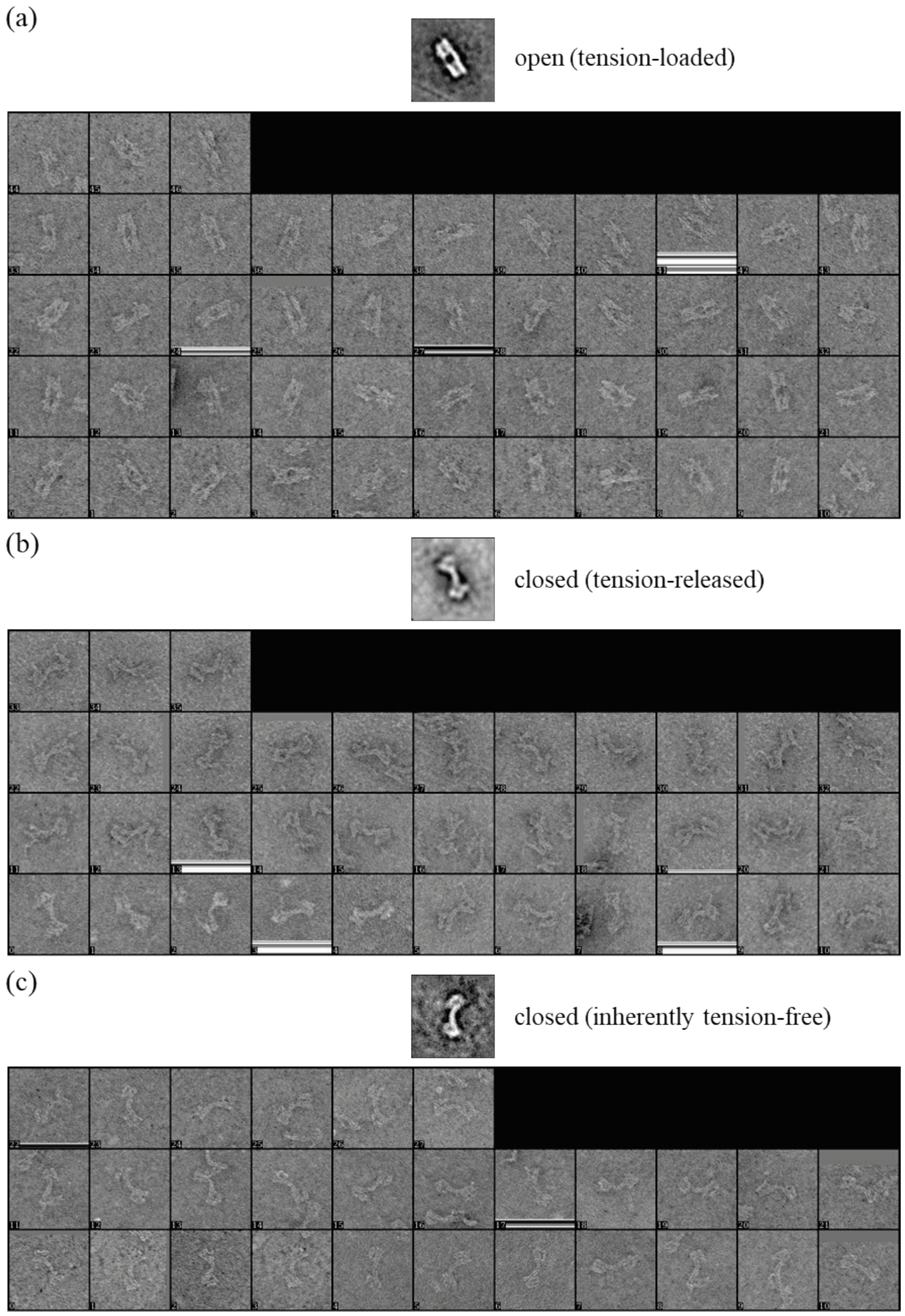

**Figure S15.** Class averaging of selected particles for DNA clamp IV. (a) open (tension-loaded) state. (b) closed (tension-released) state. (c) closed (inherently tension-free) state. Box size: 93.9 nm.

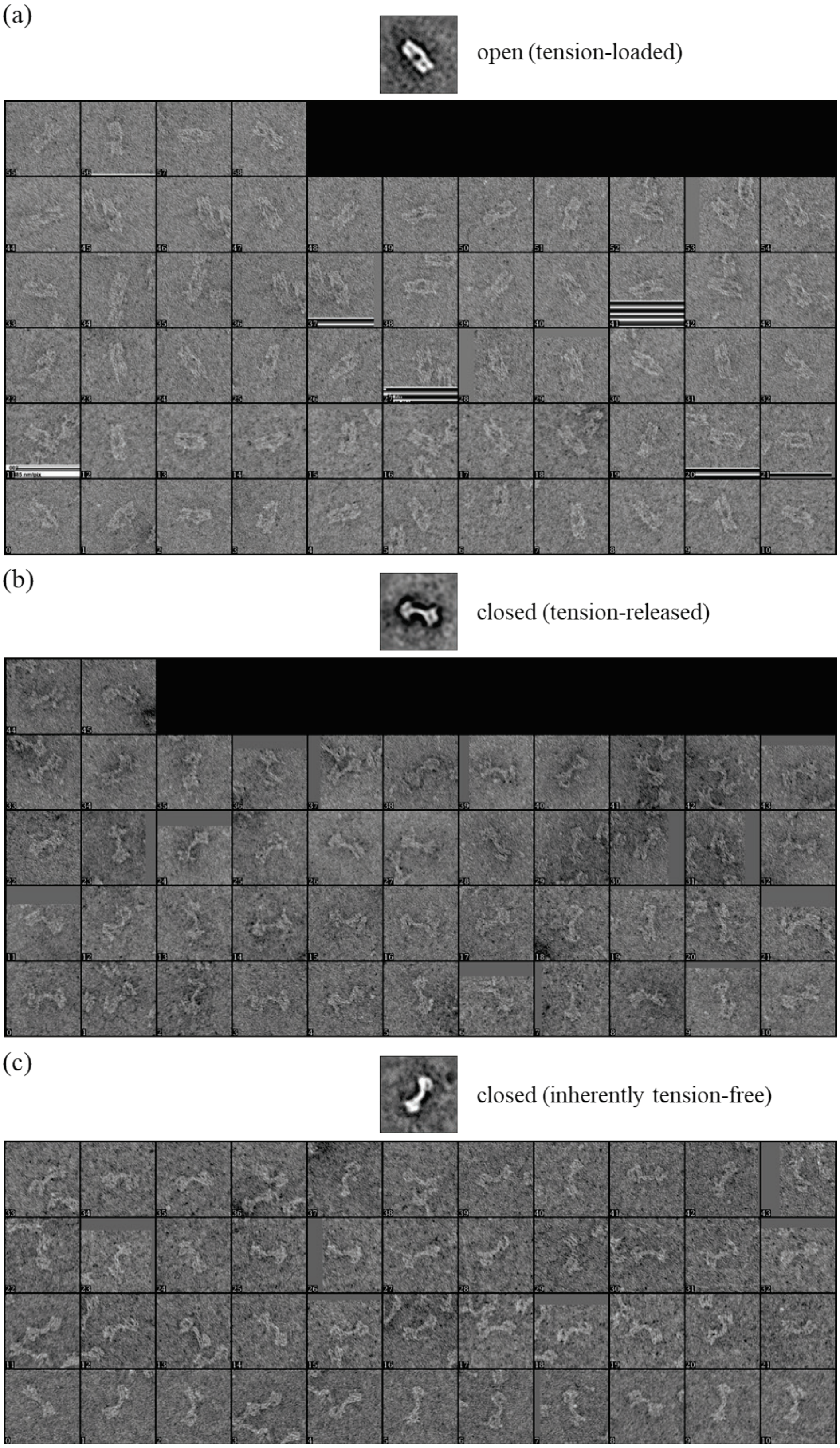

**Figure S16.** Class averaging of selected particles for DNA clamp V. (a) open (tension-loaded) state. (b) closed (tension-released) state. (c) closed (inherently tension-free) state. Box size: 93.9 nm.

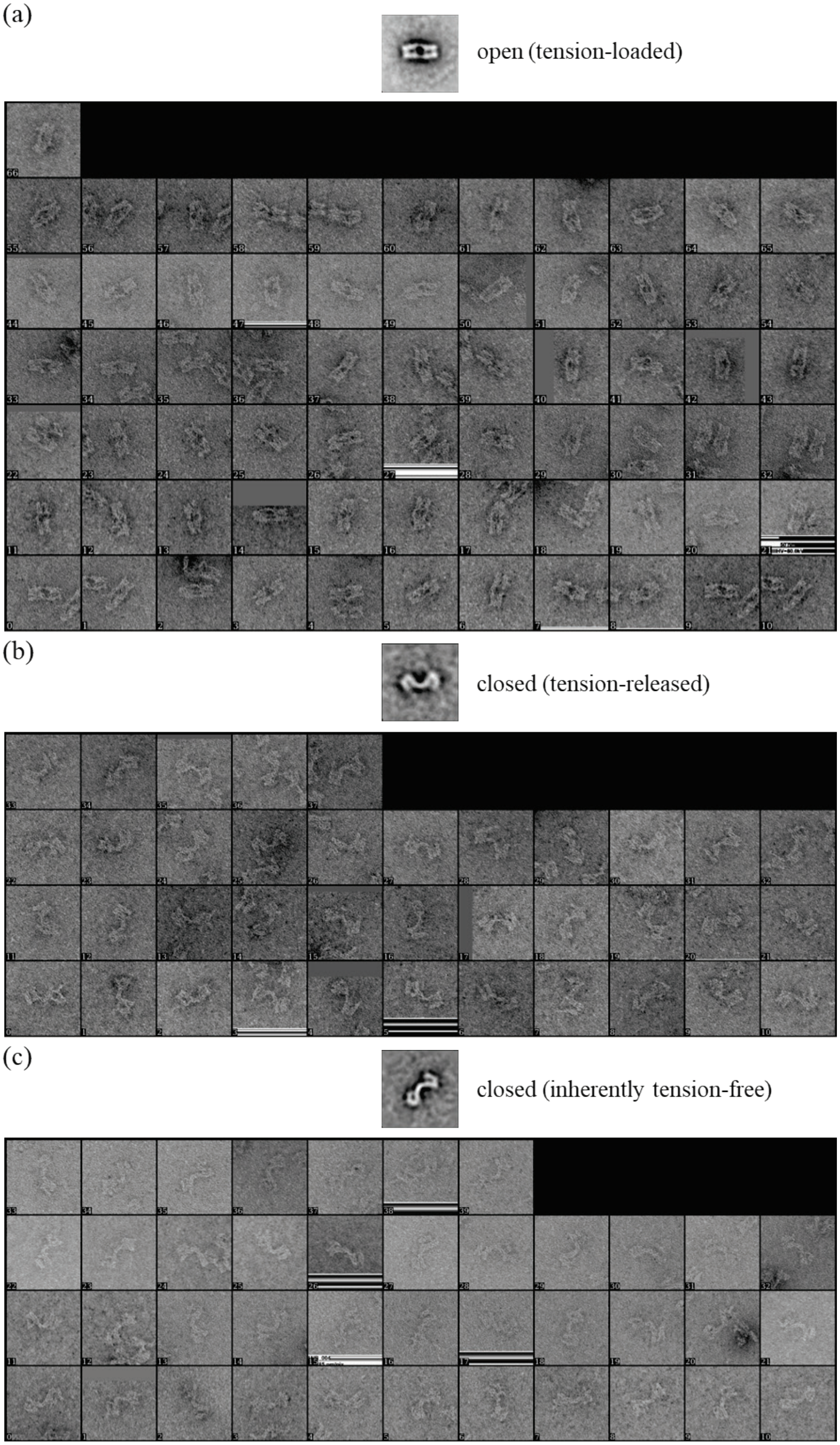

**Figure S17.** TEM images of DNA clamps outfitted with extenders and their measured bending angles (mean ± SD). Scale bars: 50 nm.

| DNA clamps | tension-loaded | tension-released | inherently tension-free |
|---|---|---|---|
| I | 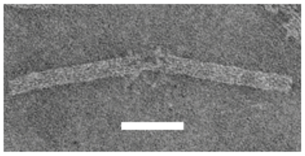 17° ± 14° | 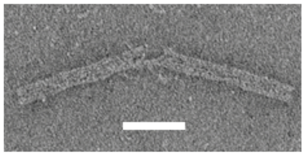 36° ± 18° | 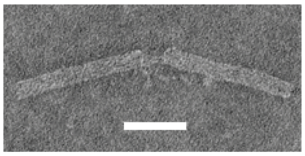 33° ± 15° |
| II | 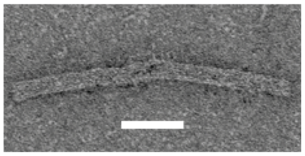 22° ± 15° | 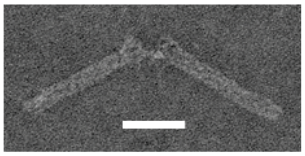 62° ± 16° | 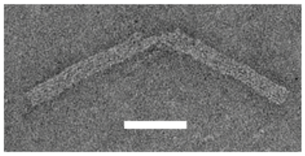 52° ± 20° |
| III | 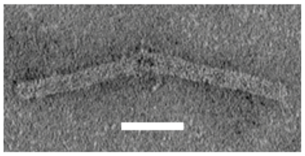 20° ± 13° | 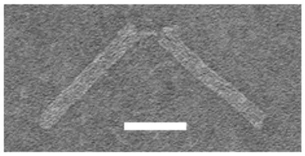 85° ± 19° | 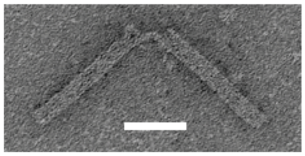 89° ± 18° |
| IV | 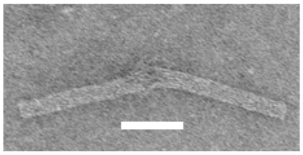 30° ± 14° | 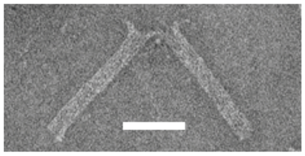 100° ± 15° | 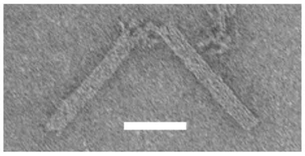 96° ± 9° |
| V | 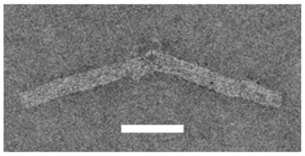 33° ± 16° | 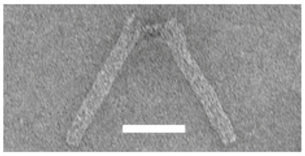 116° ± 26° | 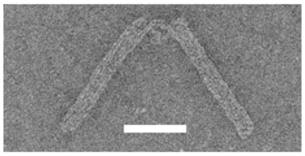 114° ± 15° |

**Figure S18.** TEM images of closed, tension-released clamps (with attached extenders for better visualization). Examples of correctly bent (red boxes) and incorrectly twisted (black boxes) clamps are shown (magnified images on the right). Ratios between correctly and incorrectly closed clamps are labeled below the magnified images. Scale bars: 100 nm. Extenders are omitted in the models on the upper right for clarity.

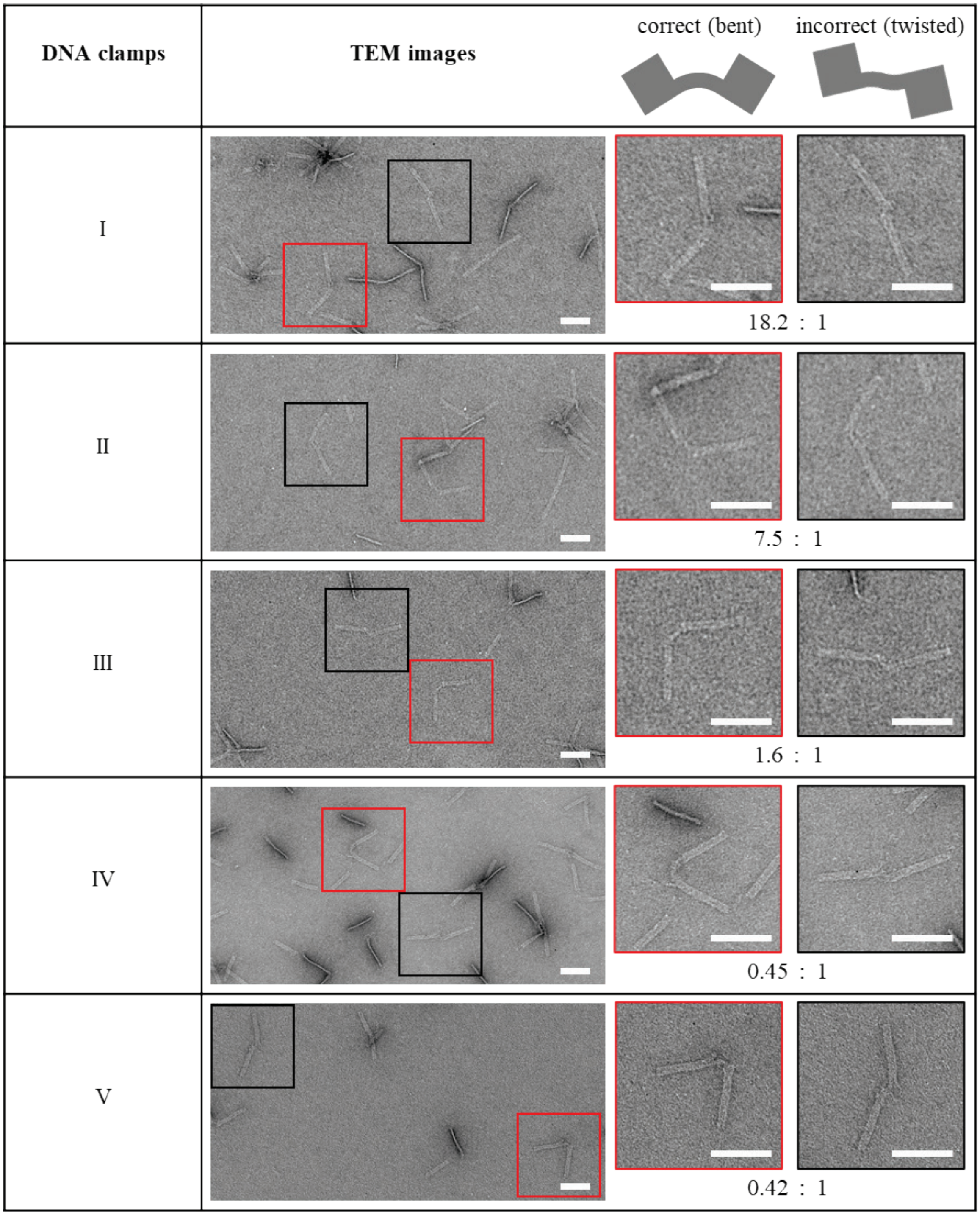

**Figure S19.** Fluorescence micrographs of DNA clamp-treated GUVs (rhod-PE: green). ITF: inherently tension-free. Scale bars: 10 μm.

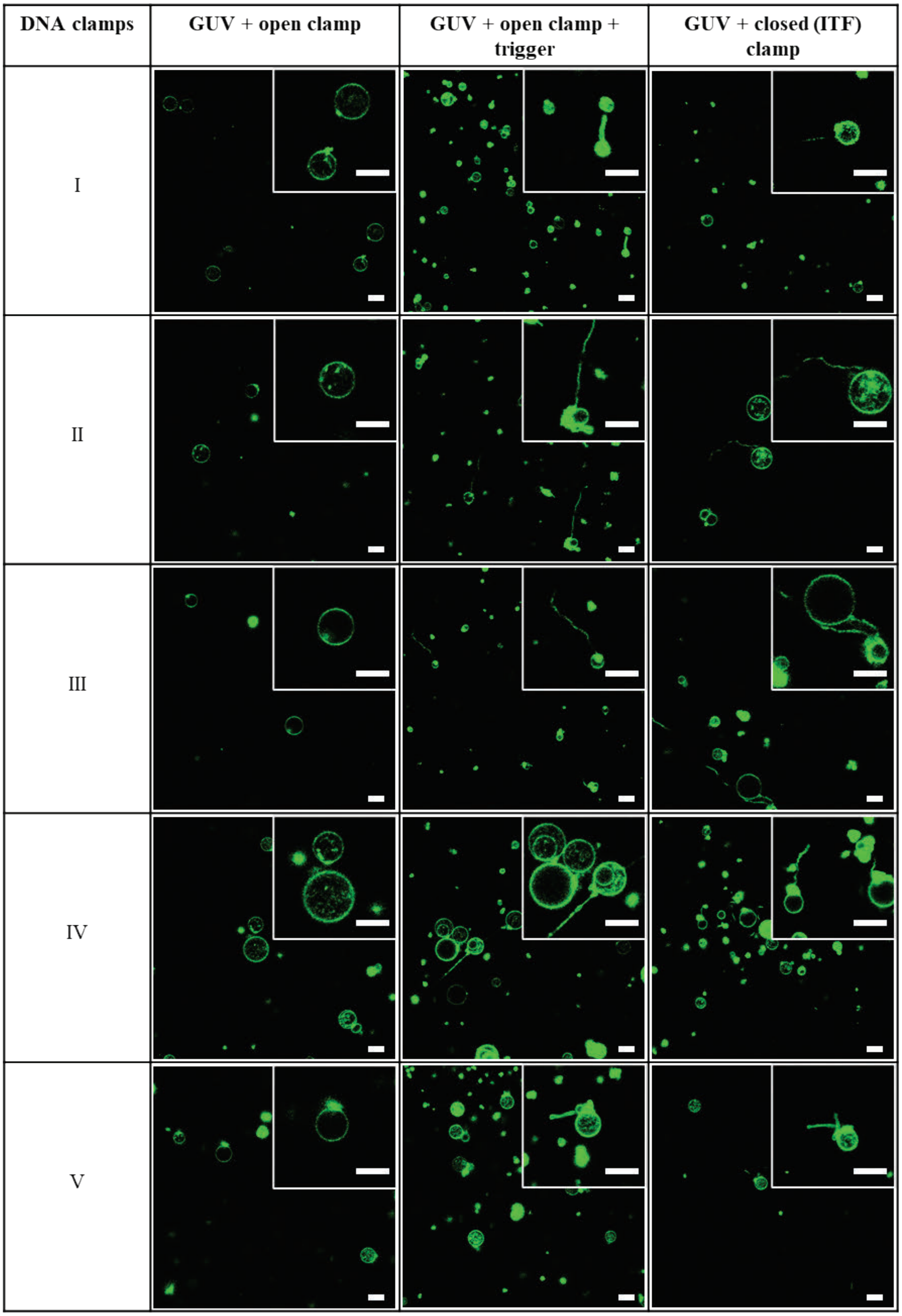

**Figure S20.** Calculation of radius of curvature of a tension-released DNA clamp ($r_c$). We assume that all cholesterols insert into the lipid bilayer and model the cross-section of cholesterol-embedded membrane as an arc (length = L, central angle = clamp bending angle = θ) of a circle (radius = $r_c$).

$$r_c = \frac{L}{\theta}$$

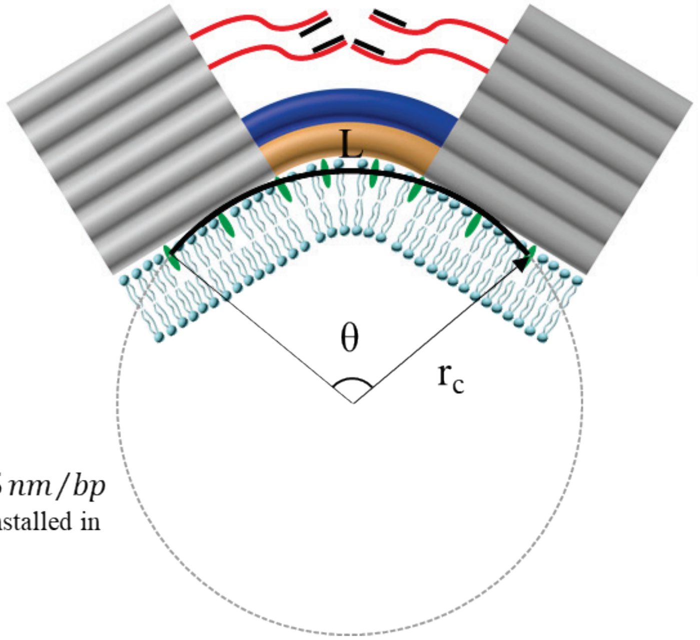


$$L = (109 - N_{indel})\ bp\ \times 0.335\, nm/bp$$

where $N_{indel}$ is the number of indel bps installed in the bridge.

**Figure S21.** TEM images of LUVs coated with DNA clamp I. ITF: inherently tension-free. Scale bars: 100 nm.

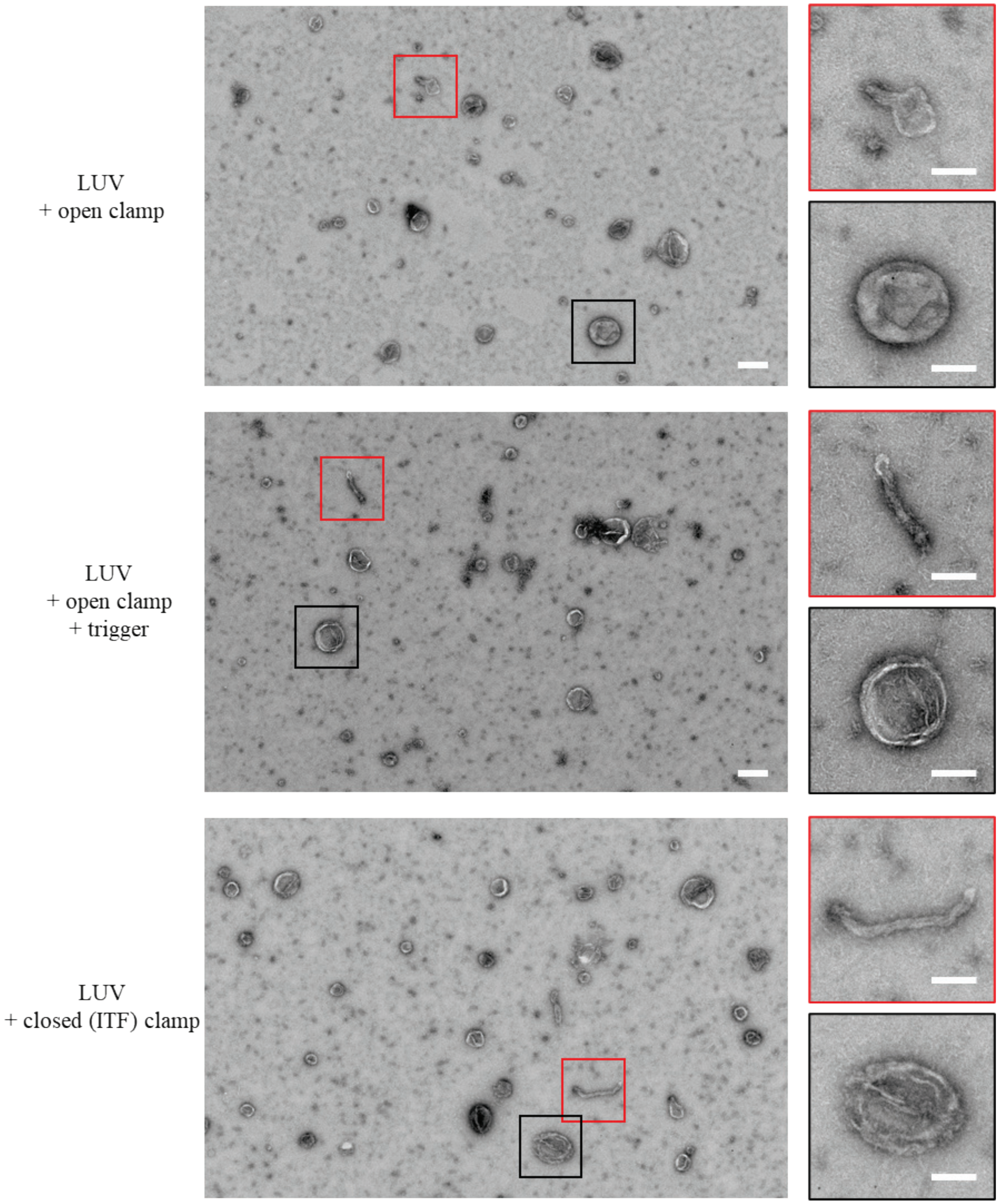

**Figure S22.** TEM images of LUVs coated with DNA clamp II. ITF: inherently tension-free. Scale bars: 100 nm.

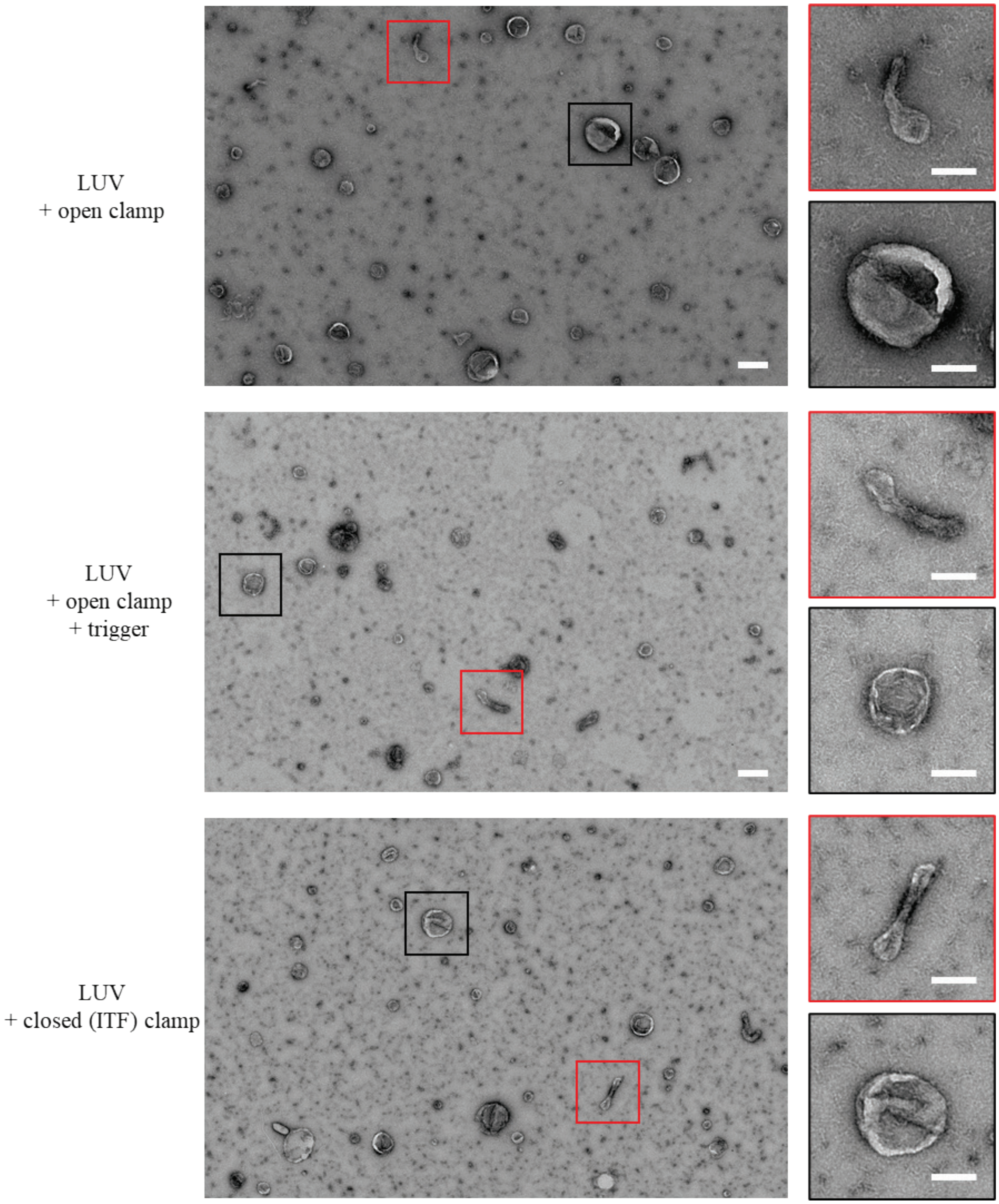

**Figure S23.** TEM images of LUVs coated with DNA clamp III. ITF: inherently tension-free. Scale bars: 100 nm.

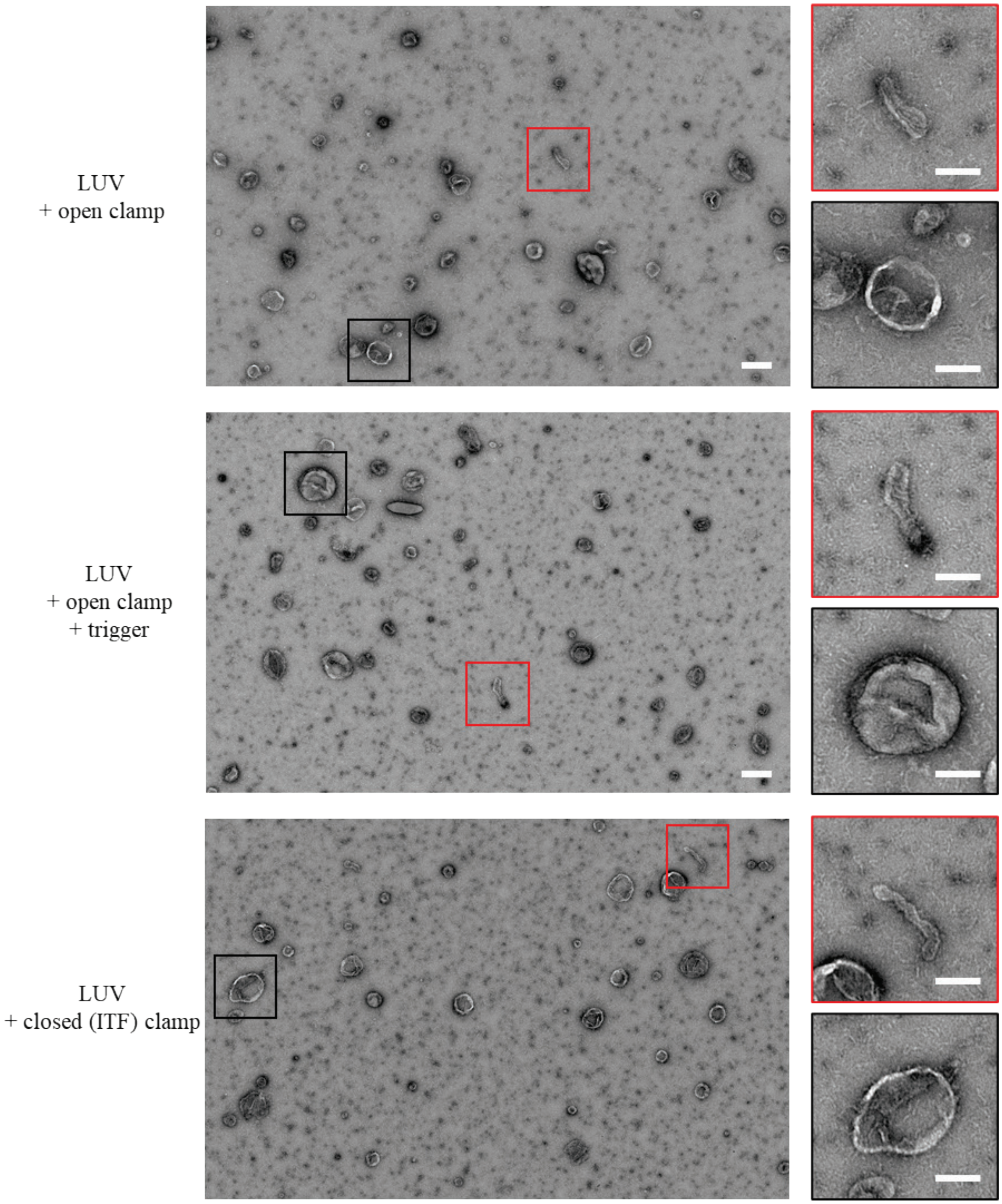

**Figure S24.** TEM images of LUVs coated with DNA clamp IV. ITF: inherently tension-free. Scale bars: 100 nm.

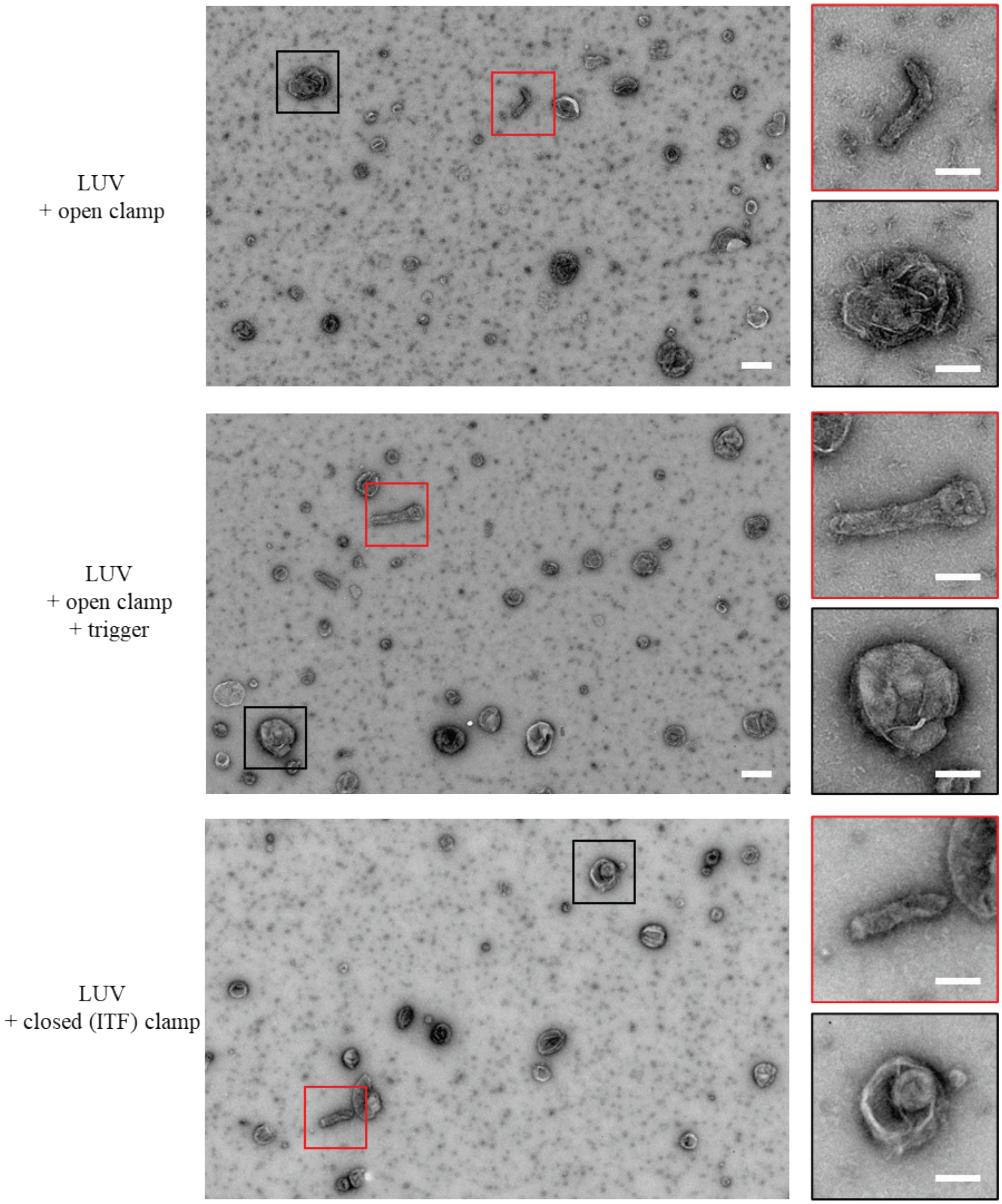

**Figure S25.** TEM images of LUVs coated with DNA clamp V. ITF: inherently tension-free. Scale bars: 100 nm.

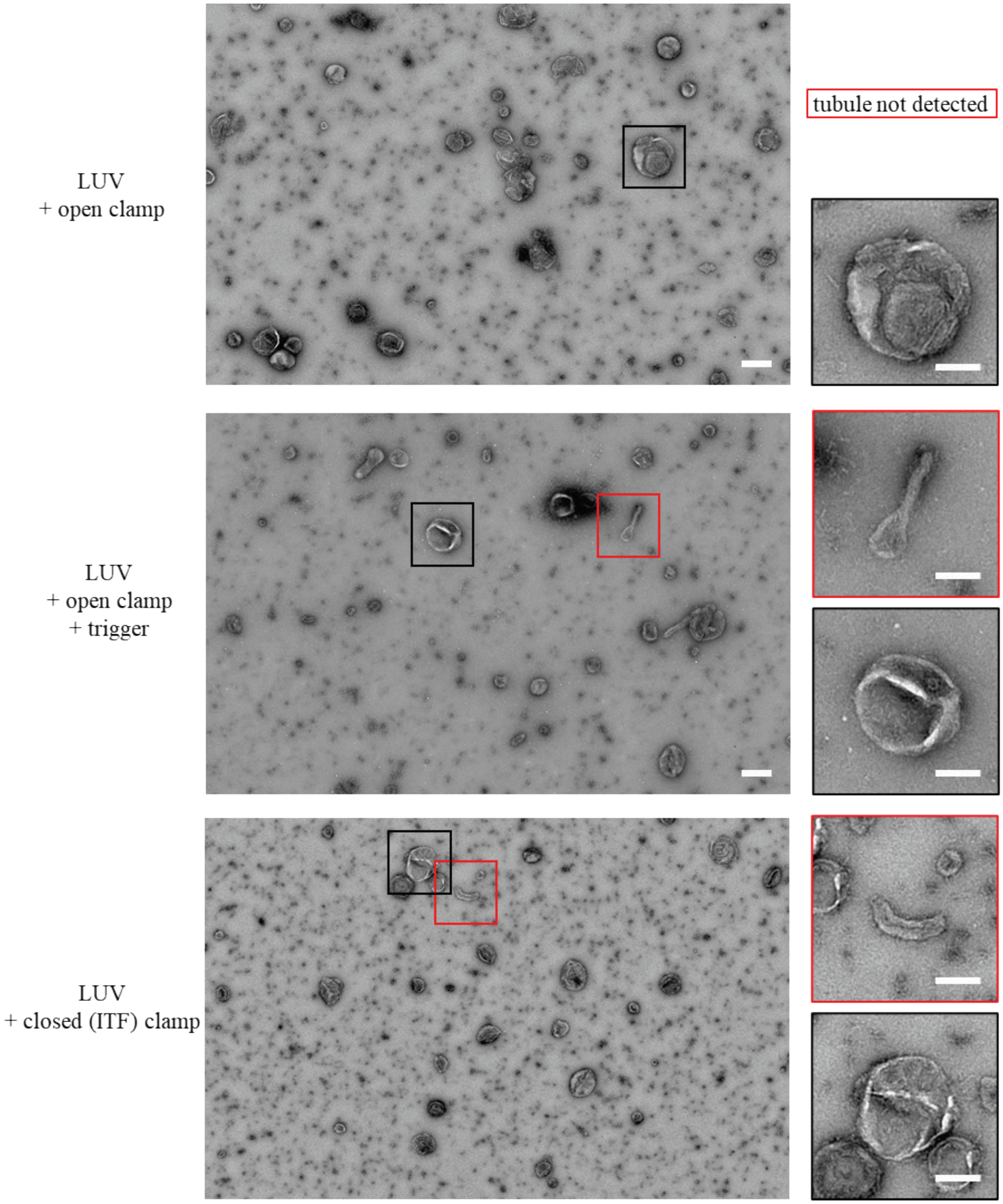

**Figure S26.** Efficiency of LUV tubulation induced by DNA clamps: (a) bar graph and (b) table. N.D.: "not detected".

(a)

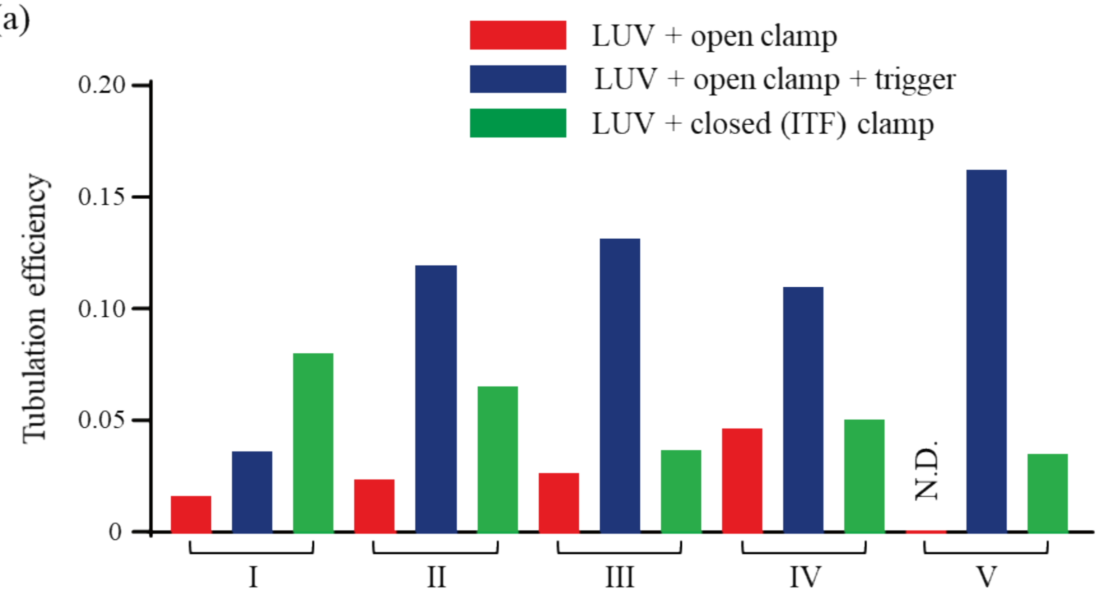


(b)

| DNA clamps | Samples | tubulation efficiency | total number of LUVs (N) |
|---|---|---|---|
| I | LUV + open clamp | 1.6% | 62 |
| | LUV + open clamp + trigger | 3.6% | 83 |
| | LUV + closed (ITF) clamp | 8.0% | 75 |
| II | LUV + open clamp | 2.3% | 86 |
| | LUV + open clamp + trigger | 11.9% | 67 |
| | LUV + closed (ITF) clamp | 6.5% | 77 |
| III | LUV + open clamp | 2.6% | 77 |
| | LUV + open clamp + trigger | 13.1% | 61 |
| | LUV + closed (ITF) clamp | 3.6% | 55 |
| IV | LUV + open clamp | 4.6% | 65 |
| | LUV + open clamp + trigger | 10.9% | 64 |
| | LUV + closed (ITF) clamp | 5.0% | 40 |
| V | LUV + open clamp | 0 | 63 |
| | LUV + open clamp + trigger | 16.2% | 37 |
| | LUV + closed (ITF) clamp | 3.5% | 57 |

### SUPPLEMENTAL TABLES

**Table S1.** Measured diameter of LUV tubules induced by actuating DNA clamps on membrane.

| DNA clamp | diameter of tubules on LUVs |
|---|---|
| I | 41.4 nm ± 11.0 nm |
| II | 36.5 nm ± 9.2 nm |
| III | 34.8 nm ± 8.3 nm |
| IV | 34.6 nm ± 12.5 nm |
| V | 30.6 nm ± 7.0 nm |

**Table S2.** DNA sequences.

| Sequence Name | Sequence |
|---|---|
| Scaffold p1512 | GGATCCACGCGCCCTGTAGCGGCGCATTAAGCGCGGC GGGTGTGGTGGTTACGCGCAGCGTGACCGCTACACTTG CCAGCGCCCTAGCGCCCGCTCCTTTCGCTTTCTTCCCTT CCTTTCTCGCCACGTTCGCCGGCTTTCCCCGTCAAGCTC TAAATCGGGGGCTCCCTTTAGGGTTCCGATTTAGTGCT TTACGGCACCTCGACCCCAAAAAACTTGATTTGGGTGA TGGTTCACGTAGTGGGCCATCGCCCTGATAGACGGTTT TTCGCCCTTTGACGTTGGAGTCCACGTTCTTTAATAGT GGACTCTTGTTCCAAACTGGAACAACACTCAACCCTAT CTCGGGCTATTCTTTTGATTTATAAGGGATTTTGCCGAT TTCGGGGTACCAACTGCTGGGCCATATCGACATGGAC ACCAGAGAATGGAGCGACGGCGTGCTCACAAACTCCG CCAGACAAGTCGTGCGCGAACCTCAAGACGTCAGCTC TTGGATCATCTGCGATGGTGATATTGACCCTGAGTGGA TCGAGTCCCTGAATTCCGTGTTGGATGACAACAGGCTC CTCACAATGCCTTCTGGTGAGAGAATCCAGTTCGGTCC TAACGTGAACTTCGTGTTCGAGACACACGATCTCAGCT GTGCTAGCCCAGCTACTATCTCCCGCATGGGAATGATC TTCTTGTCCGACGAGGAGACAGATTTGAACTCATTGAT CAAGTCTTGGCTCAGAAACCAGCCTGCAGAATATAGG AATAACCTGGAGAACTGGATCGGTGATTACTTCGAGA AGGCTTTGCAGTGGGTGCTGAAACAGAACGACTATGT CGTCGAAACCAGCCTGGTCGGTACAGTTATGAACGGA CTCTCCCATCTGCACGGATGCAGAGATCACGATGAGTT TATCATCAATTTGATCCGCGGACTGGGAGGTAACTTGA ATATGAAATCTCGCCTGGAGTTCACTAAAGAAGTGTTC CACTGGGCTAGGGAGTCACCACCTGACTTCCACAAAC CTATGGACACCTACTATGATTCCACAAGAGGCAGGTTG GCCACCTACGTGCTGAAGAAGCCTGAGGACCTCACCG CTGACGACTTCTCCAACGGACTGACTCTGCCCGTGATC CAGACTCCAGACATGCAGCGCGGACTCGATTACTTTAA GCCCTGGCTCAGCTCCGATACCAAGCAACCTTTCATTC TCGTGGGACCAGAGGGATGTGGTAAAGGAATGCTCCT GAGGTATGCATTCTCCCAGCTCCGCTCAACCCAAATTG CCACTGTTCACTGTTCAGCCCAAACAACTTCAAGGCAT CTCCTCCAGAAGCTCAGCCAGACCTGTATGGTTATCAG CACCAACACCGGCAGAGTTTACCGCCCAAAGGATTGT GAGCGCCTCGTGTTGTACCTCAAAGATATCAATCTGCC CAAACTCGATAAGTGGGGCACTTCCACCCTCGTGGCAT TTCTCCAACAGGTGCTGACCTACCAGGGCTTCTACGAC G |
| Scaffold p3024 | CCCGGTACCCAATTCGCCCTATAGTGAGTCGTATTACG CGCGCTCACTGGCCGTCGTTTTACAACGTCGTGACTGG GAAAACCCTGGCGTTACCCAACTTAATCGCCTTGCAGC ACATCCCCCTTTCGCCAGCTGGCGTAATAGCGAAGAG GCCCGCACCGATCGCCCTTCCCAACAGTTGCGCAGCCT GAATGGCGAATGGGACGCGCCCTGTAGCGGCGCATTA AGCGCGGCGGGTGTGGTGGTTACGCGCAGCGTGACCG |

| | |
|---|---|
| | CTACACTTGCCAGCGCCCTAGCGCCCGCTCCTTTCGCT<br>TTCTTCCCTTCCTTTCTCGCCACGTTCGCCGGCTTTCCC<br>CGTCAAGCTCTAAATCGGGGGCTCCCTTTAGGGTTCCG<br>ATTTAGTGCTTTACGGCACCTCGACCCCAAAAAACTTG<br>ATTAGGGTGATGGTTCACGTAGTGGGCCATCGCCCTGA<br>TAGACGGTTTTTCGCCCTTTGACGTTGGAGTCCACGTT<br>CTTTAATAGTGGACTCTTGTTCCAAACTGGAACAACAC<br>TCAACCCTATCTCGGTCTATTCTTTTGATTTATAAGGGA<br>TTTTGCCGATTTCGGCCTATTGGTTAAAAAATGAGCTG<br>ATTTAACAAAAATTTAACGCGAATTTTAACAAAATATT<br>AACGCTTACAATTTAGGTGGCACTTTTCGGGGAAATGT<br>GCGCGGAACCCCTATTTGTTTATTTTTCTAAATACATTC<br>AAATATGTATCCGCTCATGAGACAATAACCCTGATAA<br>ATGCTTCAATAATATTGAAAAAGGAAGAGTATGAGTA<br>TTCAACATTTCCGTGTCGCCCTTATTCCCTTTTTTGCGG<br>CATTTTGCCTTCCTGTTTTTGCTCACCCAGAAACGCTGG<br>TGAAAGTAAAAGATGCTGAAGATCAGTTGGGTGCACG<br>AGTGGGTTACATCGAACTGGATCTCAACAGCGGTAAG<br>ATCCTTGAGAGTTTTCGCCCCGAAGAACGTTTTCCAAT<br>GATGAGCACTTTTAAAGTTCTGCTATGTGGCGCGGTAT<br>TATCCCGTATTGACGCCGGGCAAGAGCAACTCGGTCG<br>CCGCATACACTATTCTCAGAATGACTTGGTTGAGTACT<br>CACCAGTCACAGAAAAGCATCTTACGGATGGCATGAC<br>AGTAAGAGAATTATGCAGTGCTGCCATAACCATGAGT<br>GATAACACTGCGGCCAACTTACTTCTGACAACGATCGG<br>AGGACCGAAGGAGCTAACCGCTTTTTTGCACAACATG<br>GGGGATCATGTAACTCGCCTTGATCGTTGGGAACCGG<br>AGCTGAATGAAGCCATACCAAACGACGAGCGTGACAC<br>CACGATGCCTGTAGCAATGGCAACAACGTTGCGCAAA<br>CTATTAACTGGCGAACTACTTACTCTAGCTTCCCGGCA<br>ACAATTAATAGACTGGATGGAGGCGGATAAAGTTGCA<br>GGACCACTTCTGCGCTCGGCCCTTCCGGCTGGCTGGTT<br>TATTGCTGATAAATCTGGAGCCGGTGAGCGTGGGTCTC<br>GCGGTATCATTGCAGCACTGGGGCCAGATGGTAAGCC<br>CTCCCGTATCGTAGTTATCTACACGACGGGGAGTCAGG<br>CAACTATGGATGAACGAAATAGACAGATCGCTGAGAT<br>AGGTGCCTCACTGATTAAGCATTGGTAACTGTCAGACC<br>AAGTTTACTCATATATACTTTAGATTGATTTAAAACTT<br>CATTTTTAATTTAAAAGGATCTAGGTGAAGATCCTTTT<br>TGATAATCTCATGACCAAAATCCCTTAACGTGAGTTTT<br>CGTTCCACTGAGCGTCAGACCCCGTAGAAAAGATCAA<br>AGGATCTTCTTGAGATCCTTTTTTTCTGCGCGTAATCTG<br>CTGCTTGCAAACAAAAAAACCACCGCTACCAGCGGTG<br>GTTTGTTTGCCGGATCAAGAGCTACCAACTCTTTTTCC<br>GAAGGTAACTGGCTTCAGCAGAGCGCAGATACCAAAT<br>ACTGTCCTTCTAGTGTAGCCGTAGTTAGGCCACCACTT<br>CAAGAACTCTGTAGCACCGCCTACATACCTCGCTCTGC<br>TAATCCTGTTACCAGTGGCTGCTGCCAGTGGCGATAAG<br>TCGTGTCTTACCGGGTTGGACTCAAGACGATAGTTACC<br>GGATAAGGCGCAGCGGTCGGGCTGAACGGGGGGTTCG<br>TGCACACAGCCCAGCTTGGAGCGAACGACCTACACCG |

| | |
|---|---|
| | AACTGAGATACCTACAGCGTGAGCTATGAGAAAGCGC<br>CACGCTTCCCGAAGGGAGAAAGGCGGACAGGTATCCG<br>GTAAGCGGCAGGGTCGGAACAGGAGAGCGCACGAGG<br>GAGCTTCCAGGGGGAAACGCCTGGTATCTTTATAGTCC<br>TGTCGGGTTTCGCCACCTCTGACTTGAGCGTCGATTTTT<br>GTGATGCTCGTCAGGGGGGCGGAGCCTATGGAAAAAC<br>GCCAGCAACGCGGCCTTTTTACGGTTCCTGGCCTTTTG<br>CTGGCCTTTTGCTCACATGTTCTTTCCTGCGTTATCCCC<br>TGATTCTGTGGATAACCGTATTACCGCCTTTGAGTGAG<br>CTGATACCGCTCGCCGCAGCCGAACGACCGAGCGCAG<br>CGAGTCAGTGAGCGAGGAAGCGGAAGAGCGCCCAATA<br>CGCAAACCGCCTCTCCCCGCGCGTTGGCCGATTCATTA<br>ATGCAGCTGGCACGACAGGTTTCCCGACTGGAAAGCG<br>GGCAGTGAGCGCAACGCAATTAATGTGAGTTAGCTCA<br>CTCATTAGGCACCCCAGGCTTTACACTTTATGCTTCCG<br>GCTCGTATGTTGTGTGGAATTGTGAGCGGATAACAATT<br>TCACACAGGAAACAGCTATGACCATGATTACGCCAAG<br>CGCGCAATTAACCCTCACTAAAGGGAACAAAAGCTGG<br>AGCTCCACCGCGGTGGCGGCCGCTCTAGAACTAGTGG<br>ATCCGTAAATCAATGACTTACGCGCACCGAAAGGTGC<br>GTATTGTCTATAGCCCCCTCAGCCACGAATTCGTCTGA<br>CGACGACAAGACAAGCTTGCGTGTGAATTCCCTGGCTT<br>CTCCTGAGAAA |
| cholesterol-modified anti-handle | GTGAGTTGTGGTAGATAATTT/3CholTEG/ |
| Alexa Fluor 647-labeled anti-handle | /5Alex647N/TAGATGGAGTGTGGTGTGAAG |
